# Jak2^V617F^ Reversible Activation Shows an Essential Requirement for Jak2^V617F^ in Myeloproliferative Neoplasms

**DOI:** 10.1101/2022.05.18.492332

**Authors:** Andrew Dunbar, Robert L. Bowman, Young Park, Franco Izzo, Robert M. Myers, Abdul Karzai, Won Jun Kim, Inés Fernández Maestre, Michael R. Waarts, Abbas Nazir, Wenbin Xiao, Max Brodsky, Mirko Farina, Louise Cai, Sheng F. Cai, Benjamin Wang, Wenbin An, Julie L Yang, Shoron Mowla, Shira E. Eisman, Tanmay Mishra, Remie Houston, Emily Guzzardi, Anthony R. Martinez Benitez, Aaron Viny, Richard Koche, Dan A. Landau, Ross L. Levine

## Abstract

Janus kinases (JAKs) mediate cytokine signaling, cell growth and hematopoietic differentiation.^1^ Gain-of-function mutations activating JAK2 signaling are seen in the majority of myeloproliferative neoplasm (MPN) patients, most commonly due to the *JAK2^V617F^* driver allele.^2^ While clinically-approved JAK inhibitors improve symptoms and outcomes in MPNs, remissions are rare, and mutant allele burden does not substantively change with chronic JAK inhibitor therapy in most patients.^3, 4^ This has been postulated to be due to incomplete dependence on constitutive JAK/STAT signaling, alternative signaling pathways, and/or the presence of cooperating disease alleles;^5^ however we hypothesize this is due to the inability of current JAK inhibitors to potently and specifically abrogate mutant JAK2 signaling. We therefore developed a conditionally inducible mouse model allowing for sequential activation, and then inactivation, of *Jak2^V617F^* from its endogenous locus using a Dre-*rox*/Cre-*lox* dual orthogonal recombinase system. Deletion of oncogenic *Jak2^V617F^*abrogates the MPN disease phenotype, induces mutant-specific cell loss including in hematopoietic stem/progenitor cells, and extends overall survival to an extent not observed with pharmacologic JAK inhibition. Furthermore, reversal of *Jak2^V617F^* in MPN cells with antecedent loss of *Tet2*^6, 7^ abrogates the MPN phenotype and inhibits mutant stem cell persistence suggesting cooperating epigenetic-modifying alleles do not alter dependence on mutant JAK/STAT signaling. Our results suggest that mutant-specific inhibition of JAK2^V617F^ represents the best therapeutic approach for JAK2^V617F^-mutant MPN and demonstrate the therapeutic relevance of a dual-recombinase system to assess mutant-specific oncogenic dependencies *in vivo*.

## MAIN

Somatic mutations which constitutively activate JAK2 signaling are seen in the majority of MPN patients,^2^ most commonly the recurrent *JAK2^V617F^* alteration, and murine models suggest a critical role for JAK/STAT pathway mutations in promoting the MPN phenotype *in vivo*.^8–12^ In contrast to ABL1 kinase inhibition in BCR-ABL1-driven chronic myelogenous leukemia,^13^ current JAK inhibitors fail to reduce mutant clonal fraction, do not induce pathologic regression of key disease features including myeloproliferation and bone marrow fibrosis, and most patients lose their response over time.^3, 4^ To date, second-site JAK2 mutations have not been observed as a mechanism of acquired resistance,^14^ and different mechanisms have been postulated to mediate the inadequate efficacy of JAK inhibition, including incomplete dependency on JAK2 signaling and the presence of co-occurring mutant disease alleles.^15^ We hypothesized that the limited potency of JAK inhibition relates to insufficient mutant kinase inhibition at achievable therapeutic doses, and we and others have elucidated mechanisms by which mutant JAK2 can signal in the presence of type I JAK inhibitors.^16–18^ However, no study to date has shown that potent, specific inhibition of mutant JAK2 can induce disease regressions, and current model systems do not accurately recapitulate reversal of endogenous mutant expression or allow assessment of oncogenic dependency in the clonal evolution to myeloid transformation. Given this, we developed a system which would definitively assess JAK2^V617F^ dependency in MPN.

### A conditional knock-in, knock-out model of *Jak2^V617F^* MPN

To assess dependency of MPN maintenance on Jak^V617F^ oncogenic signaling, we generated a Dre- *rox*,^19^ Cre-*lox*^20^ dual-recombinase *Jak2^V617F^* knock-in/knock-out mouse model (*Jak2^Rox/Lox^*/*Jak2^RL^*) by gene targeting in mouse embryonic stem cells **(Fig. 1a)**. The close proximity of the *lox* sites (82 base pairs) prevents Cre-mediated deletion prior to Dre-mediated recombination and *Jak2^V617F^*induction. Once the mutant allele is activated, the *lox* sites separate allowing for subsequent Cre-mediated deletion of *Jak2^V617F^*, including in models where cooperating alleles are induced by antecedent Cre-mediated activation/deletion. A similar strategy, which we have termed GOLDI- Lox for <u>g</u>overning <u>o</u>ncogenic loci by <u>D</u>re inversion and *<u>lox</u>* deletion (GOLDI-Lox), was used to target *Flt3^ITD^* in the accompanying manuscript (see Bowman, R. *et al.* 2022). Sequencing of the *Jak2^RL^* locus on sorted Cre reporter cells^21^ after Cre recombinase exposure confirmed retainment of the non-recombined *Jak2^RL^* locus **(Extended Data Fig. 1a)**. We used mRNA electroporation of Dre mRNA *ex vivo* to transiently express the Dre recombinase in primary hematopoietic stem/progenitor cells (HSPCs), which efficiently induces *Jak2^V617F^* activation and separation of *lox* sites by inversion **(Extended Data Fig. 1b-c)**. By three weeks post-transplant, lethally irradiated mice transplanted with Dre-inducible *Jak2^RL^* knock-in bone marrow developed a highly penetrant and fully transplantable MPN characterized by leukocytosis with myeloid preponderance, elevated hematocrit with erythroid progenitor expansion in bone marrow, hepatosplenomegaly, and megakaryocytic hyperplasia consistent with prior *Jak2^V617F^* conditional knock-in mouse models of MPN **(Extended Data Fig. 1d-h)**.^9^ Progressive bone marrow fibrosis was also observed at 24 weeks in 9 of 15 *Jak2^RL^* knock-in mice.

**Figure 1:**
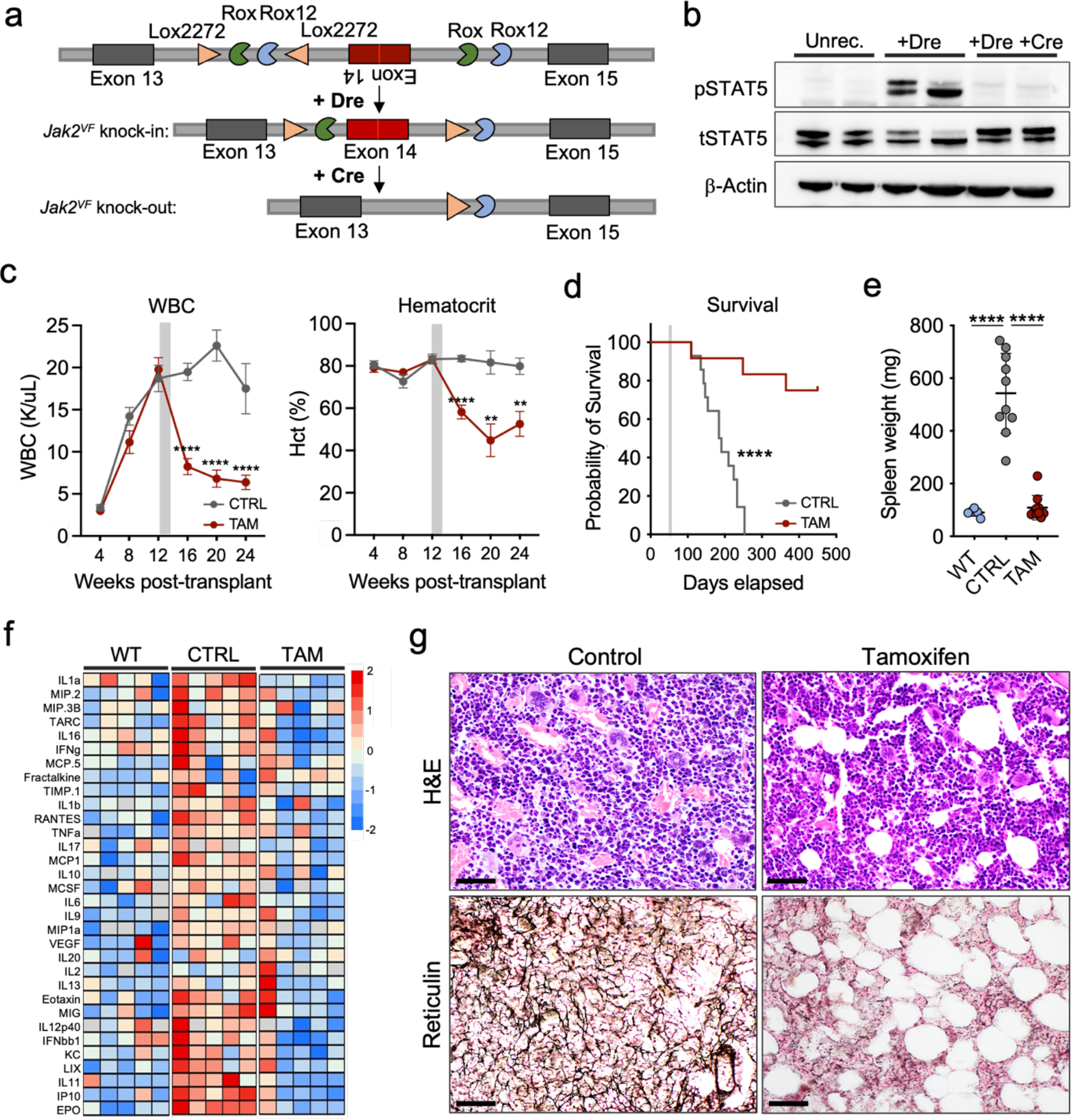
*Jak2^V617F^* deletion abolishes JAK/STAT signaling and abrogates the MPN phenotype. **a.** Schematic representation of the dual-recombinase *Jak2^V617F^*conditional knock- in/knock-out allele (*Jak2^RL^*), the *Jak2^RL^* knock-in allele following Dre recombination, and the null recombined allele following Cre-mediated deletion. Semi-circles indicate *Rox* sequences; triangles indicate *lox*P sequences. **b.** Representative western blot depicting phospho-STAT5 abundance of Dre-mediated *Jak2^V617F^* knock-in (+Dre) vs. *Jak2^V617F^-*deleted (+Dre +Cre) states from isolated splenocytes 7 days following tamoxifen administration in comparison to unrecombined (Unrec.) *Jak2^RL^* cells (*n* = 2 biological replicates each; representative of *n* = 2 independent experiments). **c.** Peripheral blood count trends (weeks 0-24) of MPN (CTRL) vs. tamoxifen (TAM; *Jak2^V617F^*- deleted) treated mice: white blood cells (WBC; left panel), hematocrit (Hct; right panel) (*n* ≥ 10 per arm; mean ± s.e.m). Gray bar represents duration of TAM pulse/chow administration. Representative of *n* = 2 independent transplants. \*\**p* ≤ 0.01, \*\*\*\**p* ≤ 0.0001. **d.** Kaplan–Meier survival analysis of MPN (CTRL) vs. tamoxifen (TAM; *Jak2^V617F^*-deleted) treated mice (*n* ≥ 12 per arm; Log-rank test). Gray bar represents duration of TAM pulse/chow administration. \*\*\*\**p* ≤ 0.0001. **e.** Spleen weights of MPN (CTRL) vs. tamoxifen (TAM; *Jak2^V617F^*-deleted) treated mice at timed sacrifice (24 weeks) in comparison to wild-type control mice (mean ± s.e.m.). Representative of *n* = 2 independent transplants. \*\*\*\**p* ≤ 0.0001. **f.** Heatmap demonstrating cytokine/chemokine concentrations (log10 concentration; pg/mL) in peripheral blood serum of MPN (CTRL) vs. tamoxifen (TAM; *Jak2^V617F^*-deleted) treated mice in comparison to wild-type control mice (*n* = 5 biological replicates per arm). **g.** Representative hematoxylin and eosin (H&E) and reticulin stains of bone marrow of MPN (Control) vs. tamoxifen (*Jak2*^V617F^-deleted) treated mice from timed sacrifice 24 weeks. Representative micrographs of *n* = 6 individual mouse replicates per arm. All images represented at 400X magnification. Scale bar: 20μm.

To assess the reversibility of the *Jak2^RL^*construct, we cultured Dre-electroporated, lineage-negative, tamoxifen-inducible Ubc:CreER-*Jak2^RL^* cells isolated from donor mice with active MPN *ex vivo* with increasing doses of 4-hydroxy-tamoxifen (4-OHT) over bone marrow endothelial cells (BMECs) **(Extended Data Fig. 2a)**.^22^ Treatment with 4-OHT resulted in deletion of the *Jak2^V617F^* allele, which was confirmed by excision polymerase chain reaction (PCR) **(Extended Data Fig. 2b)**. Loss of *Jak2^V617F^* significantly reduced cell numbers *ex vivo* (mean 0.18 × 10^6^/mL vs. 2.19 × 10^6^/mL, *p* ≤ 0.0001), including within immunophenotypically-defined HSPC compartments, a phenotypic change not observed with vehicle-treated *Jak2^RL^*, Cre-inducible *Jak2^V617F^*(*Jak2^Crelox^*; *p* ≤ 0.228),^8^ or Cre-inducible wild-type (WT) cells (*p* ≤ 0.114) **(Extended Data Fig. 2c-g)**. Loss of *Jak2^V617F^* also abrogated erythropoietin-independent erythroid differentiation^23^ *in vitro* (*p* ≤ 0.01), an effect not observed when cells were exposed to the JAK inhibitor ruxolitinib^24^ **(Extended Data Fig. 2h)**. The cell loss observed was associated with enhanced apoptosis, which was most apparent in Mac1^+^ mature myeloid cells (mean 35% vs. 9.3%, *p* ≤ 0.005) (**Extended Data Fig. 2i)**.

**Figure 2:**
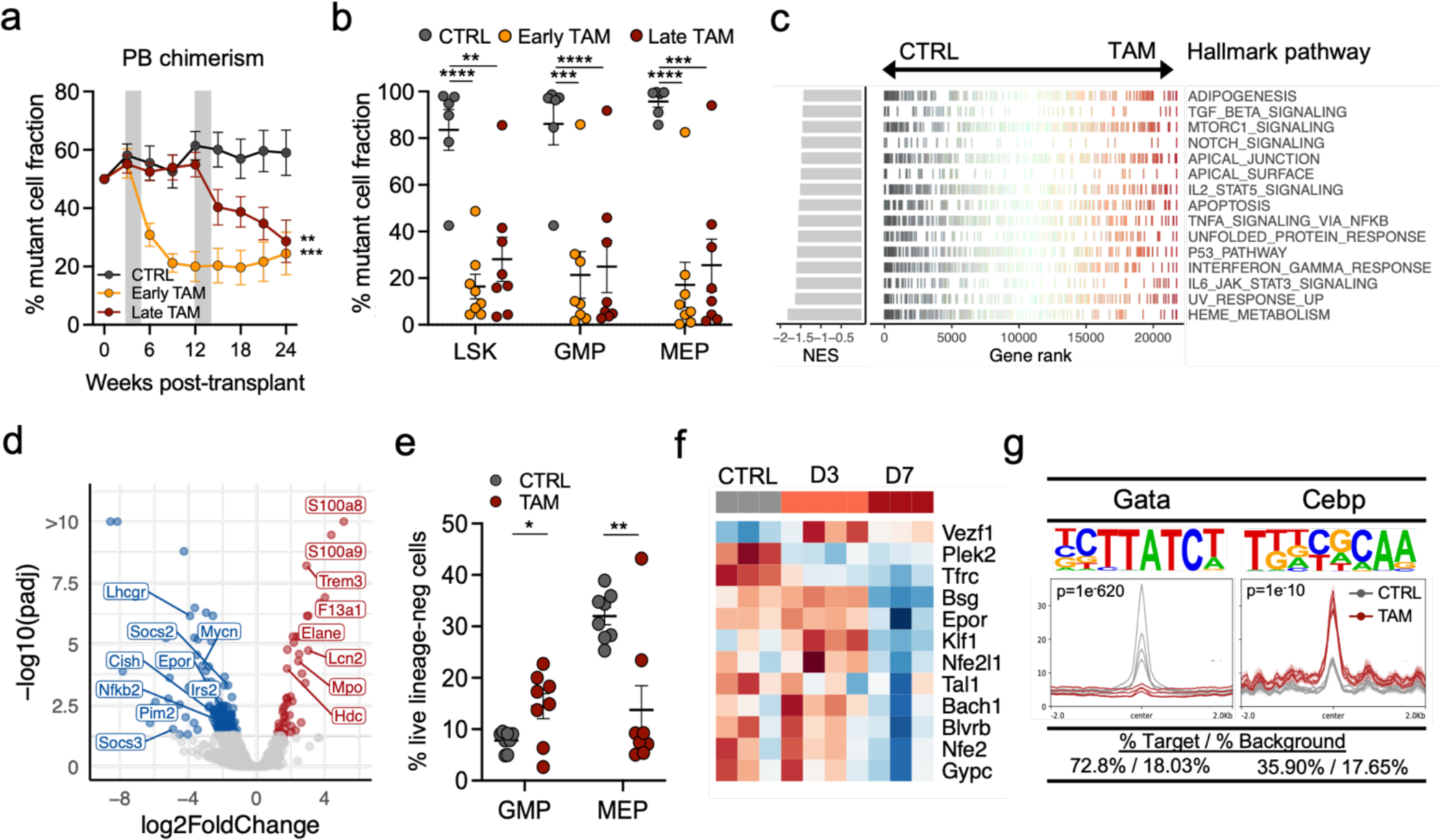
*Jak2^V617F^* reversal impairs the fitness of MPN cells, including MPN stem cells. **a.** Peripheral blood (PB) mutant Cd45.2 percent chimerism trend (weeks 0-24) of early (3 weeks post-transplant) tamoxifen (TAM; *Jak2^V617F^*-deleted) treated (gold bar) and late (12 weeks post- transplant) tamoxifen treated (maroon bar) mice (*n* = 8 each) in comparison to MPN (CTRL; dark gray bar; *n* = 6) mice (mean ± s.e.m.). Gray bars represent duration of TAM pulse/chow administration. Representative of *n* = 2 independent transplants. \*\**p* ≤ 0.01, \*\*\**p* ≤ 0.001. **b.** Bone marrow mutant cell fraction within LSK (Lineage^-^Sca1^+^cKit^+^), granulocytic-monocytic progenitor (GMP; Lineage^-^cKit^+^Sca1^-^Cd34^+^Fcg^+^), and megakaryocytic-erythroid progenitor (MEP; Lineage^-^ cKit^+^Sca1^-^Cd34^-^Fcg^-^) compartments of early (3 weeks post-transplant) tamoxifen (TAM; *Jak2^V617F^*-deleted) treated and late (12 weeks post-transplant) tamoxifen treated mice in comparison to MPN (CTRL) mice at timed sacrifice 24 weeks (*n* = 6-8 individual biological replicates per arm; mean ± s.e.m.). Representative of *n* = 2 independent transplants. \*\**p* ≤ 0.01, \*\*\**p* ≤ 0.001, \*\*\*\**p* ≤ 0.0001. **c.** Gene-set enrichment analysis (GSEA) of significant Hallmark gene sets of MPN (CTRL) vs. tamoxifen (TAM; *Jak2^V617F^*-deleted) treated LSKs isolated 7 days following initiation of TAM (*n* = 3-4 biological replicates per arm). **d.** Volcano plot demonstrating differential gene expression of MPN (CTRL) vs. TAM (*Jak2^V617F^*-deleted) treated LSKs 7 days following initiation of TAM (*n* = 3-4 biological replicates per arm). **e.** GMP and MEP stem cell frequencies of MPN (CTRL) vs. tamoxifen (TAM; *Jak2^V617F^*-deleted) treated mice 7 days following initiation of TAM (*n* = 8 biological replicates per arm across 2 independent transplants; mean ± s.e.m.). **f.** Row normalized heatmap of RNA-sequencing data of key erythroid differentiation factor genes from harvested MEPs at baseline (CTRL), day 3 (D3) and day 7 (D7) following initiation of TAM (*Jak2^V617F^* deletion). **g.** HOMER motif analysis from ATAC-seq data demonstrating decreased accessibility of Gata motif signatures with concomitant increased accessibility of Cebp motif signatures of TAM treated (*Jak2^V617F^*-deleted) cKit^+^ bone marrow cells isolated 7 days following initiation of treatment in comparison to MPN (CTRL) cells (*n* = 3 biological replicates per arm).

We next evaluated the impact of reversible *Jak2^V617F^*expression *in vivo*. Twelve weeks post-transplant, secondary recipient mice transplanted with Dre-electroporated Ubc:CreER-*Jak2^RL^*whole bone marrow and exhibiting MPN were administered tamoxifen (TAM) to delete *Jak2^V617F^* **(Extended Data Fig. 3a)**. A sequential dual recombinase reporter system^25^ was used to validate *Jak2^V617F^* deletion within Cd45.2 reporter-positive cell populations **(Extended Data Fig. 3b)**. Deletion of *Jak2^V617F^* was also validated *in vivo* at the transcriptional level (*p* ≤ 0.0001) **(Extended Data Fig. 3c)** and was associated with loss of constitutive JAK/STAT signaling **(Fig. 1b)**. Consistent with our *in vitro* data, we observed normalization of white blood cell (WBC; mean 6.18 K/uL vs. 17.5 K/uL, *p* ≤ 0.0001), hematocrit (Hct; mean 52.6% vs. 79.9%, p ≤ 0.01), and platelet (mean 786 K/uL vs. 2146 K/uL, *p* ≤ 0.0004) parameters within 4 weeks following tamoxifen treatment that persisted until timed sacrifice at 24 weeks **(Fig. 1c, Extended Data Fig. 3d)**. Two of 12 mice demonstrated reemergence/persistence of the MPN phenotype, both of which showed incomplete excision of the *Jak2^RL^* allele highlighting the necessity of Jak2^V617F^ in disease maintenance **(Extended Data Fig. 3e)**. Genetic reversal of *Jak2^V617F^* significantly prolonged overall survival (median not defined vs. 187 days, *p* ≤ 0.0012) and led to loss of disease-defining MPN features in the majority of mice (9/12) **(Fig. 1d)**. Spleen weights (mean 108.9 mg vs. 542.7 mg, *p* ≤ 0.0001) and inflammatory cytokine levels were also reduced to normal levels in mice with *Jak2^V617F^* reversal **(Fig. 1e-f)**. Histopathologic analysis revealed reductions in megakaryocytic hyperplasia, splenic infiltration, and absence of bone marrow and spleen fibrosis in 8 of 9 assayed *Jak2^V617F^*-deleted mice that persisted until timed sacrifice at 24 weeks **(Fig. 1g, Extended Data Fig 3f)**. The phenotypes observed with Ubc:CreER-*Jak2^V617F^* deletion *in vivo* were not observed with tamoxifen administration in the absence of *Jak2^V617F^*reversal **(Extended Data Fig. 4)**. We conclude that the MPN phenotype requires maintenance of oncogenic signaling through Jak2^V617F^.

**Figure 3:**
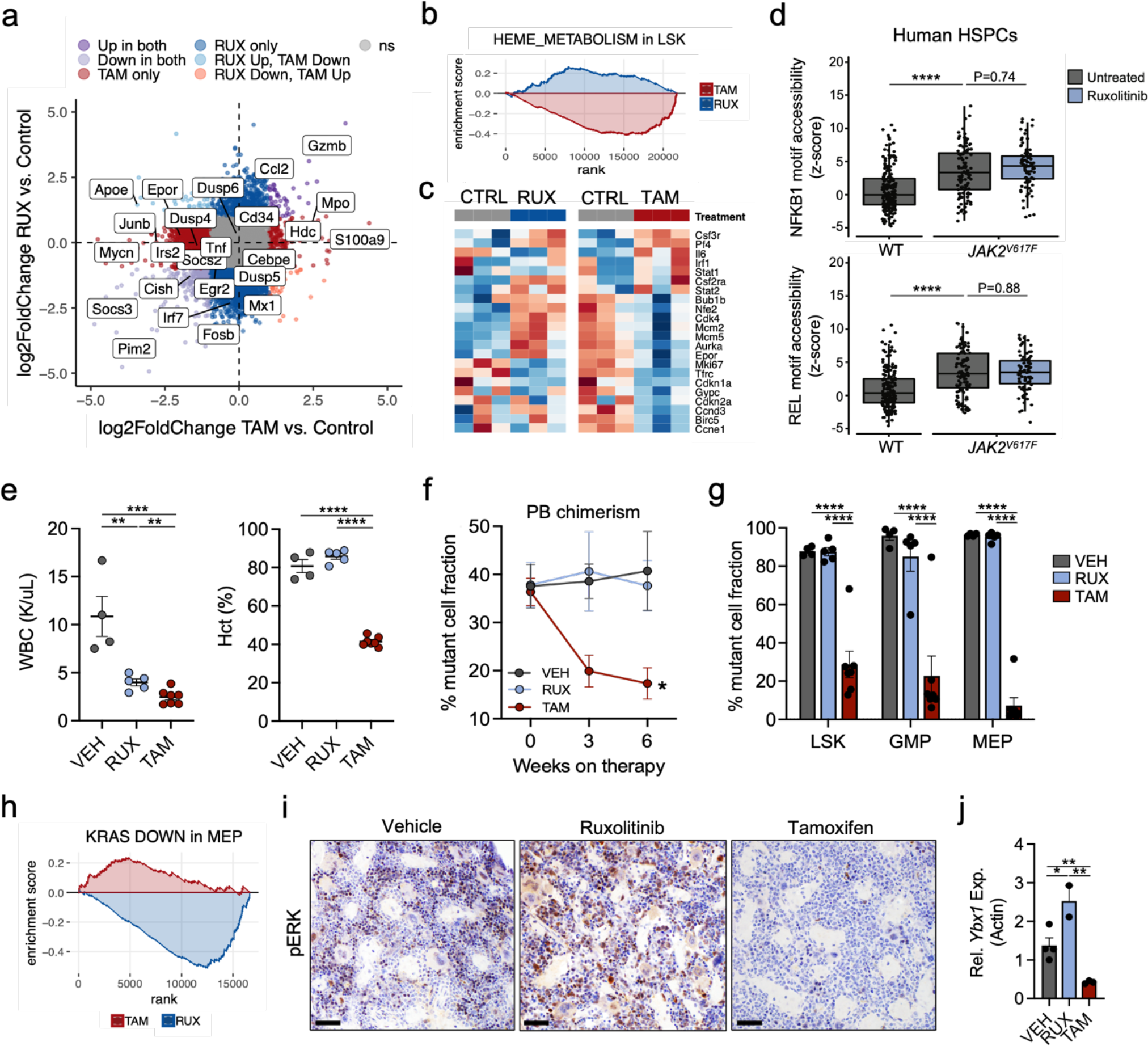
Differential efficacy of *Jak2^V617F^* deletion compared to JAK inhibitor therapy. **a.** Scatter plot depicting log2FoldChange of ruxolitinib (RUX) treated vs. tamoxifen (TAM; *Jak2^V617F^*-deleted) treated LSKs (Lineage^-^Sca1^+^cKit^+^) in comparison to MPN control LSKs isolated after 7 days of treatment (*n* = 2-3 biological replicates per arm); differentially expressed genes as indicated by color (see Supplemental Tables 1 and 3). **b.** Gene set enrichment analysis (GSEA) depicting a positive enrichment in heme metabolism in ruxolitinib (RUX) treated (*n* = 3) vs. negative enrichment in tamoxifen (TAM; *Jak2^V617F^*-deleted) treated (*n* = 3) LSKs isolated after 7 days of treatment. **c.** Row normalized heatmap of RNA-sequencing data of ruxolitinib (RUX) treated (blue) or tamoxifen (TAM; *Jak2^V617F^*-deleted) treated (red) megakaryocytic-erythroid progenitor (MEP; Lineage^-^cKit^+^Sca1^-^Cd34^-^Fcg^-^) cells in comparison to MPN (CTRL) cohorts (gray). **d.** Box plots of single-cell ATAC-seq motif accessibility for either NFKB1 or REL transcription factors for untreated human *JAK2* wild-type (*n* = 188 cells from 4 patients; gray), untreated *JAK2^V617F^*-mutant (*n* = 105 cells from 4 patients; gray), and ruxolitinib- treated *JAK2^V617F^*-mutant (*n* = 87 cells from 3 patients; blue) HSPCs (from Myers, R.M. and Izzo, F. *et al*., bioRxiv, 2022). *P* values indicated are from linear mixture model explicitly modeling patient identity as random effect to account for patient-specific effects, followed by likelihood ratio test. \*\*\*\**p* ≤ 0.0001. **e.** Peripheral blood counts of vehicle (VEH), ruxolitinib (RUX), or tamoxifen (TAM; *Jak2^V617F^*-deleted) treated mice at the conclusion of a 6-week *in vivo* trial: white blood cells (WBC; left panel), hematocrit (Hct; right panel) (*n* ≥ 4 each; mean ± s.e.m). \*\**p* ≤ 0.01, \*\*\**p* ≤ 0.001, \*\*\*\**p* ≤ 0.0001. **f.** Peripheral blood (PB) mutant Cd45.2 percent chimerism trend (0-6 weeks) of vehicle (VEH), ruxolitinib (RUX), or tamoxifen (TAM; *Jak2^V617F^*-deleted) treated mice (*n* ≥ 4 each; mean ± s.e.m.). \**p* ≤ 0.05. **g.** Bone marrow mutant cell fraction of LSK (Lineage^-^ Sca1^+^cKit^+^), granulocytic-monocytic progenitor (GMP; Lineage^-^cKit^+^Sca1^-^Cd34^+^Fcg^+^), and megakaryocytic-erythroid progenitor (MEP; Lineage^-^cKit^+^Sca1^-^Cd34^-^Fcg^-^) compartments of vehicle (VEH), ruxolitinib (RUX), or tamoxifen (TAM; *Jak2^V617F^*-deleted) treated mice at the conclusion of the 6-week *in vivo* trial (*n* ≥ 4 each; mean ± s.e.m). \**p* ≤ 0.05, \*\*\*\**p* ≤ 0.0001. **h.** GSEA depicting a negative enrichment in down-regulation of KRAS signaling targets in ruxolitinib (RUX) treated (*n* = 3) vs. positive enrichment in tamoxifen (TAM; *Jak2^V617F^*-deleted) treated (*n* = 3) MEPs isolated following 7 days of respective treatment. **i.** Immunohistochemistry of phospho-ERK on sectioned bone marrow of vehicle, ruxolitinib, or tamoxifen (*Jak2^V617F^*- deleted) treated mice following 7 days of treatment (*n* = 3 individual biological replicates per arm). All images represented at 400X magnification. Scale bar: 20μm. **j.** Quantitative polymerase-chain reaction demonstrating relative *Ybx1* expression levels from isolated cKit^+^ bone marrow of vehicle (VEH) vs. ruxolitinib (RUX) vs. tamoxifen (TAM; *Jak2^V617F^*-deleted) treated mice following 7 days of treatment (*n* = 2-4 individual biological replicates per arm; mean ± s.e.m). \**p* ≤ 0.05, \*\**p* ≤ 0.01. **e-g.** Representative of *n* = 3 independent experiments.

**Figure 4:**
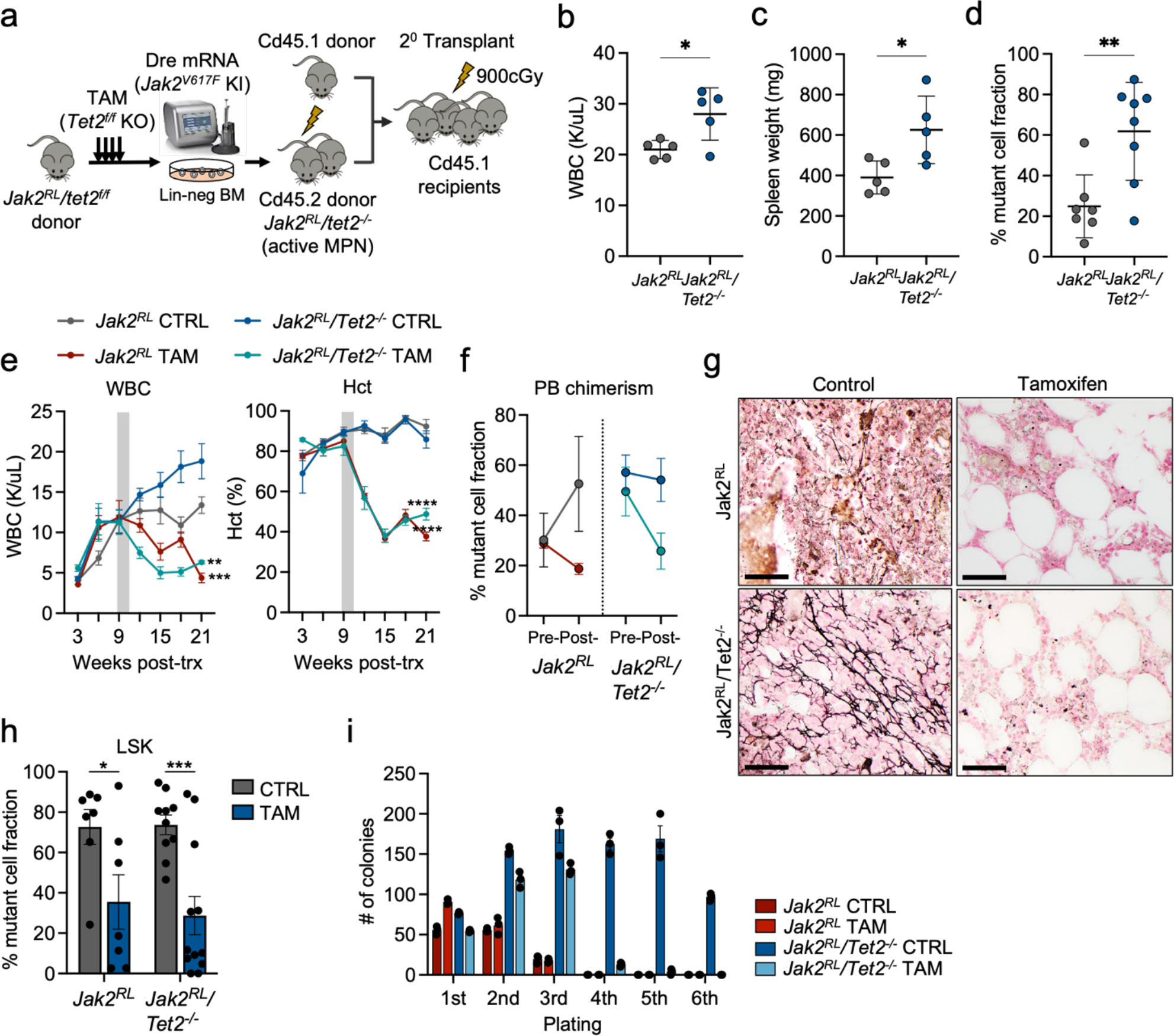
Jak2^V617F^ dependency with cooperative Tet2 loss. **a.** Schematic of the experimental set up for the double-mutant *Jak2^RL^/Tet2^f/f^*competitive transplants. TAM: tamoxifen; KI: knock- in; KO: knock-out; Lin-neg BM: lineage-negative bone marrow; cGy: centigray. Downward arrows represent initial pulse TAM administration to genetically inactivate *Tet2*. **b.** White blood cell (WBC) counts of primary *Jak2^RL^* vs. *Jak2^RL^/Tet2^-/-^* transplanted mice at 16 weeks post- transplant (*n* = 5 each; mean ± s.d.). Representative of an *n* = 2 independent transplants. \**p* ≤ 0.05. **c.** Spleen weights of primary *Jak2^RL^* vs. *Jak2^RL^/Tet2^-/-^* transplanted mice at time of sacrifice (*n* = 5 each; mean ± s.d.). Representative of an *n* = 2 independent transplants. \**p* ≤ 0.05. **d.** Peripheral blood Cd45.2 mutant percent chimerism of *Jak2^RL^* vs. *Jak2^RL^/Tet2^-/-^*secondary competitive transplant mice at 16 weeks post-transplant (*n =* 7-8 each; mean ± s.d.). Representative of an *n* = 2 independent transplants. \*\**p* ≤ 0.01. **e.** Peripheral blood count trends (weeks 0-21) of MPN (CTRL) vs. tamoxifen (TAM; *Jak2^V617F^*-deleted) treated *Jak2^RL^* vs. *Jak2^RL^/Tet2^-/-^* competitive transplant mice: white blood cells (WBC; left panel), hematocrit (Hct; right panel) (*n* = 3-4 per arm; mean ± s.e.m). Gray bars represent duration of TAM pulse/chow administration. Representative of *n* = 2 independent transplants. \*\**p* ≤ 0.01, \*\*\**p* ≤ 0.001, \*\*\*\**p* ≤ 0.0001. **f.** Percent change in Cd45.2 mutant peripheral blood chimerism pre- vs. post-tamoxifen (*Jak2^V617F^*- deletion) treatment of *Jak2^RL^* vs. *Jak2^RL^/Tet2^-/-^* mice in relation to Cd45.1 competitor cells (*n* = 3- 4 per arm; mean ± s.e.m). Representative of *n* = 2 independent transplants. **g.** Reticulin stains of bone marrow from MPN control vs. tamoxifen (*Jak2*^V617F^-deleted) treated *Jak2^RL^* vs. *Jak2^RL^/Tet2^- /-^* mice at timed sacrifice 21 weeks. Representative micrographs of *n* = 3 individual mouse replicates per arm. All images represented at 400X magnification. Scale bar: 20μm. **h.** Bone marrow mutant Cd45.2 percent chimerism within the LSK (Lineage^-^Sca1^+^cKit^+^) compartment of MPN (CTRL) vs. tamoxifen (TAM; *Jak2*^V617F^-deleted) treated *Jak2^RL^* vs. *Jak2^RL^/Tet2^-/-^* mice at timed sacrifice 21 weeks (*n* ≥ 7 biological replicates per arm across 2 independent transplants; mean ± s.e.m). \**p* ≤ 0.05, \*\*\**p* ≤ 0.001. **i.** Serial replating assay of plated MPN (CTRL) vs. tamoxifen (TAM; *Jak2*^V617F^-deleted) treated *Jak2^RL^* vs. *Jak2^RL^/Tet2^-/-^* bone marrow cells harvested at timed sacrifice 21 weeks and scored at day 8 after each plating (each sample plated in triplicate, representative of *n* = 2 independent experiments, mean ± s.d.).

### *Jak2^V617F^* reversal impairs the fitness of MPN cells, including MPN stem cells

We next evaluated Dre-electroporated *Jak2^RL^*bone marrow from Cd45.2 MPN donors in competition with Cd45.1 competitor cells to explore effects of *Jak2^V617F^* deletion on peripheral blood and bone marrow mutant cell fitness **(Extended Data Fig. 5a)**. We observed abrupt, durable reductions in Cd45.2 mutant cell fraction in the peripheral blood (mean 24.5% vs. 63.9%, *p* ≤ 0.001) with *Jak2^V617F^*reversion that coincided with normalization of hematologic parameters and persisted until time of sacrifice, either with early (3 weeks) or late (12 weeks) post-transplant administration of tamoxifen **(Fig. 2a, Extended Data Fig. 5b)**. Consistent with the *in vitro* data, this effect was most pronounced in Mac1^+^ myeloid cell fractions (*p* ≤ 0.0001) **(Extended Data Fig. 5c)**. In bone marrow, the reduction in mutant cell fraction among different HSPC compartments was more pronounced, including within megakaryocytic-erythroid progenitor (MEP; Lineage^-^cKit^+^Sca1^-^Cd34^-^Fcg^-^; *p* ≤ 0.0001) and granulocytic-monocytic progenitor (GMP; Lineage^-^cKit^+^Sca1^-^Cd34^+^Fcg^+^; *p* ≤ 0.0001) populations and down to the level of the LSK (Lineage^-^ Sca1^+^cKit^+^; *p* ≤ 0.0096) stem cell compartment, including primitive long-term hematopoietic stem cells (LT-HSCs; Lineage^-^Sca1^+^cKit^+^Cd150^+^Cd48^-^) **(Fig. 2b, Extended Data Fig. 5d-e)**. Recurrent MPN, as was seen in the non-competitive setting, was observed in 3 of 14 mice across both early- and late- treatment arms and corresponded with residual mutant Jak2^V617F^ expression and sustained mutant chimerism in remaining mice at sacrifice. Transplant of unfractionated *Jak2^RL^*-deleted bone marrow failed to form phenotypic disease in 4 of 5 secondary transplant recipient mice consistent with depletion of disease-propagating MPN stem cells **(Extended Data Fig. 5f-h)**.

We sought to characterize the transcriptional changes seen when *Jak2^V617F^* is acutely reversed. We performed RNA sequencing (RNA-Seq) analysis of purified HSPCs 3 and 7 days following *Jak2^V617F^* deletion (*n* = 3-4) compared to MPN controls (*n* = 3-4). Transcriptional analysis of sorted, *Jak2^V617F^*-deleted LSK and MEP populations revealed near-complete loss of expression of STAT5 target genes as early as 3 days post-deletion (LSK: NES -1.77, *FDR* ≤ 0.002; MEP: NES -1.53, *FDR* ≤ 0.0065) indicating immediate disengagement from disease-defining pathway signaling **(Extended Data Fig. 6a)**. By 7 days, we observed significant negative enrichment in IFNγ (NES -1.61, *FDR* ≤ 0.0005), TGF≤ (NES -1.45, *FDR* ≤ 0.071), and TNFα via NFκB (NES - 1.54, *FDR* ≤ 0.0017) Hallmark pro-inflammatory response pathways, as well as down-regulation of MAPK (NES -1.52, *FDR* ≤ 0.0052) and MTORC1 (NES -1.46, *FDR* ≤ 0.0071) targets in LSKs suggesting abrupt reduction in pro-inflammatory and proliferative signaling in the setting of *Jak2^V617F^* deletion **(Fig. 2c, Extended Data Fig. 6b, Supplemental Table 1).** A flux towards increased expression of myeloid genes sets compared to erythroid gene sets was also observed, characterized by increased *S100a8*, *S100a9*, *Mpo*, and *Hdc* expression in LSKs, increases in GMP (mean 14.5% vs. 7.8%, *p* ≤ 0.018) vs. MEP (mean 13.8% vs. 32%, *p* ≤ 0.0025) frequencies within the HSPC compartment, and enrichment in bone marrow Mac1^+^ myeloid cells (mean 41.7% vs. 27.8%, *p* ≤ 0.0084) (**Fig. 2d-e, Extended Data Fig. 6c)**. In line with reduced erythroid output, we also observed a marked decrease in heme metabolism in MEPs (NES -2.07, *FDR* ≤ 4.71x10^-5^) with associated reductions in critical erythroid/megakaryocytic transcription factors and signaling mediators, including *Nfe3*,^26, 27^ *Plek2*,^28^ and *EpoR*^29^ which coincided with concomitant reductions in total erythroid progenitor cell numbers (*p* ≤ 0.021) and significantly reduced burst forming unit-erythroid (BFU-E) colony output of *Jak2^V617F^* deleted cells (*p* ≤ 0.001) **(Fig. 2f, Extended Data Fig. 6d-f)**. Assay for Transposase Accessible Chromatin with high-throughput sequencing (ATAC-seq) on *Jak2^V617F^*-deleted cKit^+^ cells demonstrated an increase in open chromatin with Cebp motifs (*p* ≤ 1x10^-10^) and reduced accessibility at Gata motifs (*p* ≤ 1x10^-620^), including at critical erythroid loci (e.g. *EpoR*; log2FC 1.49, *FDR* ≤ 0.00135), further consistent with an erythroid to myeloid lineage switch **(Fig. 2g, Extended Data Fig. 6g, Supplemental Table 2)**. While reduced accessibility at putative Gata target sites was observed, we did not observe differential expression of either *Gata1* (*p* ≤ 1.0) or *Gata2* (*p* ≤ 0.82) in *Jak2^V617F^*-deleted LSKs or MEPs compared to controls. These data suggest the transcriptional networks regulating the MPN phenotype are not obligately achieved through transcription factor expression dysregulation but through differential transcription factor-mediated output.

### Differential efficacy of *Jak2^V617F^* deletion compared to JAK inhibitor therapy

Given the limited ability of current JAK inhibitors to achieve disease modification and/or clonal remissions in polycythemia vera and myelofibrosis, we next compared the phenotypic and transcriptional effects of JAK inhibitor therapy with ruxolitinib to the effects of *Jak2^V617F^* reversal. We first performed RNA-seq on *Jak2^V617F^*-mutant LSKs and MEPs following 7 days of ruxolitinib treatment (*n* = 3) and compared this to the effects of *Jak2^V617F^* deletion (*n* = 3). Analysis revealed that JAK-STAT target gene expression and erythroid pathway gene expression were much less potently inhibited with ruxolitinib than with *Jak2^V617F^* deletion. Specifically, *Jak2^V617F^* deletion resulted in a significant reduction in expression of negative regulators of JAK/STAT signaling (NES -1.51, *p* ≤ 0.003) including *Socs2*,^30^ *Pim2*,^31^ and *Cish*^32^ **(Fig. 3a, Extended Data Fig. 7a, Supplemental Table 3)**. Meanwhile, ruxolitinib treatment was associated with a muted reduction in the same targets **(Extended Data Fig. 7a)**, with no significant changes in STAT5 target gene expression identified by GSEA (NES -0.913, *p* = 0.84). Furthermore, the alterations in erythroid pathway gene expression (NES 1.45, *p* ≤ 0.012 vs. NES -1.82, *p* ≤ 0.0005) and skewing of myeloid/erythroid progenitor frequencies observed with *Jak2^V617F^* deletion were not observed with ruxolitinib (mean GMP: vehicle [VEH] 6.93% vs. ruxolitinib [RUX] 6.66% vs. TAM 20.1%, *p* = 0.91 vs. *p* ≤ 0.0001, MEP: VEH 27.1% vs. RUX 35.3% vs. TAM 14.2%, *p* = 0.25 vs. *p* = 0.014)

**(Fig. 3b-c, Extended Data Fig. 7b)**. Expression of the gene sets associated with TGF≤ (*p* = 0.65) and TNFα/NFκB (*p* = 0.90) inflammatory signaling pathways were also not significantly altered with ruxolitinib treatment, in contrast to *Jak2^V617F^* deletion. Consistent with this lack of change, genotype-aware single-cell ATAC-Seq (scATAC-Seq) on myelofibrosis (MF) patient samples demonstrated unaltered NFκB accessibility in *JAK2^V617F^*-mutant HSPCs following JAK inhibitor treatment **(Fig. 3d, Extended Data Fig. 7c;** see Myers, R. and Izzo, F. *et al*., bioRxiv, 2022) supporting the notion of insufficient mitigation of inflammatory signaling by JAK inhibition on MPN-sustaining stem cells.

To evaluate the phenotypic effects of *Jak2^V617F^*deletion in direct comparison to JAK kinase inhibition, we performed an *in vivo* trial lasting 6 weeks comparing ruxolitinib to *Jak2^V617F^* deletion **(Extended Data Fig. 7d)**. We saw a greater improvement in hematologic parameters, spleen weights (mean VEH 457 mg vs. RUX 235 mg vs. TAM 125 mg, *p* ≤ 0.0027), restoration of histopathologic morphology in both bone marrow and spleen, and reduced Cd45.2 mutant chimerism in peripheral blood (mean VEH 40.7% vs. RUX 37.7% vs. TAM 17.3%, *p* ≤ 0.0059) of *Jak2^V617F^*-deleted mice versus ruxolitinib treated mice **(Fig. 3e-f, Extended Data Fig. 7e-g)**. Most importantly, the reduction in mutant cell fraction seen with *Jak2^V617F^* deletion within hematopoietic progenitor (GMP: *p* ≤ 0.0001, MEP: *p* ≤ 0.0001) and LSK stem cell enriched populations (mean VEH 87.9% vs. RUX 87.6% vs. TAM 28.7%, *p* ≤ 0.0001) was not observed with pharmacologic JAK inhibition **(Fig. 3g)**. These data indicate that current JAK inhibitors do not sufficiently inhibit mutant JAK2 signaling and that more potent target inhibition offers the potential for greater therapeutic efficacy.

Previous studies have suggested that MAPK signaling plays an important role in MPN disease cell survival in the setting of type I JAK inhibitor therapy,^17^ and recent work has implicated the MAPK- dependent factor YBX1 as a critical mediator of *JAK2^V617F^*-mutant cell persistence.^18^ We observed distinct effects on MAPK activity by RNA-Seq with ruxolitinib treatment vs. *Jak2^V617F^* deletion in comparison to vehicle treated mice. Negative regulators of KRAS signaling were down-regulated with ruxolitinib (NES -1.64, *FDR* ≤ 0.0005) and up-regulated with *Jak2^V617F^* deletion (NES 1.35, *FDR* ≤ 0.039) suggesting enhanced MAPK signaling with ruxolitinib and MAPK attenuation with *Jak2^V617F^* deletion **(Fig. 3h)**. Immunohistochemistry of bone marrow sections confirmed increased phospho-ERK abundance in ruxolitinib-treated mice that was abrogated with *Jak2^V617F^* deletion **(Fig. 3i)**, and genotype-specific scATAC-seq revealed increased accessibility of MAPK-mediated AP-1 factors FOS/JUN^33^ within HSPCs of ruxolitinib-treated MF patients in comparison to untreated MF HSPCs consistent with enhanced MAPK activity **(Extended Data Fig. 7h)**. Furthermore, expression of *Ybx1* in sorted murine cKit^+^ cells was increased with ruxolitinib therapy but potently suppressed with *Jak2^V617F^* deletion (mean rel. exp. VEH 1.37 vs. RUX 2.52 vs. TAM 0.42, *p* ≤ 0.0094) **(Fig. 3j)**. These data suggest that potent, mutant-specific *Jak2^V617F^*targeting can abrogate pathologic MAPK signaling and Ybx1-mediated persistence of *Jak2^V617F^*- mutant HSPCs.

### Jak2^V617F^ dependency with cooperative Tet2 loss

Previous studies of mutational order in primary MPN cells have shown that cooperating mutations in epigenetic regulators, including TET2, can precede the acquisition of JAK2^V617F^ in the clonal evolution of MPN and that antecedent TET2 mutations can alter the *in vitro* sensitivity to ruxolitinib.^34^ In addition, *in vitro* and *in vivo* studies have shown that concurrent TET2 and JAK2^V617F^ mutations promote enhanced mutant hematopoietic stem cell (HSC) fitness and increased risk of disease progression.^35–37^ Our *Jak2^RL^* system allows for the assessment of Jak2^V617F^ dependency in the setting of co-occurring mutant allele activation/inactivation, including in the context of antecedent mutations in epigenetic regulators. We therefore assessed the impact of *Jak2^V617F^*activation in concert with pre-existing *Tet2* loss with the reversible *Jak2^RL^* allele **(Fig. 4a)**. Mice transplanted with Dre-electroporated Ubc:CreER*-Jak2^RL^/Tet2^-/-^* cells demonstrated enhanced leukocytosis (mean 27.0 K/μL vs. 21.0 K/μL, *p* ≤ 0.021), increased spleen volumes (mean 626 mg vs. 374.8 mg, *p* ≤ 0.0047), expanded mutant peripheral blood chimerism (mean 61.9% vs. 24.9%, *p* ≤ 0.0042), and improved serial replating capacity in colony forming assays compared to Ubc:CreER*-Jak2^RL^*transplanted mice **(Fig. 4b-d, Extended Fig. 8a-b)**. These data are consistent with previous *Jak2^V617F^/Tet2^-/-^*models^35, 36^ and highlight the utility of the Dre-Cre dual recombinase system to model evolution from pre-malignant, clonally restricted hematopoietic states (i.e. clonal hematopoiesis [CH]) to overt MPN. *Ex vivo* co-culture of *Jak2^RL^/Tet2^-/-^* cells over BMECs exhibited a near 2-fold increase in hematopoietic cell output (mean 2.32 × 10^6^/mL vs.

1.18 × 10^6^/mL, *p* 0.0017), particularly among Mac1^+^ mature myeloid cells (mean 0.69 × 10^6^/mL vs. 0.35 × 10^6^/mL, *p* 0.011), compared to *Jak2^RL^* cells consistent with the known role of *Tet2* loss-of-function in enhancing myeloid lineage commitment **(Extended Data Fig. 8c)**.^6^

We next evaluated effects of *Jak2^V617F^*deletion on *Jak2^RL^/Tet2^-/-^* mutant cell fitness *in vivo* in competition with Cd45.1 bone marrow. Treatment with tamoxifen at 9 weeks post-transplant resulted in normalization of hematologic parameters (*p* ≤ 0.005) and reductions in peripheral blood mutant cell fraction of double-mutant cells to a similar extent observed with *Jak2^V617F^* deletion in single-mutant *Jak2^RL^* transplanted mice **(Fig. 4e-f)**. Further, spleen sizes (mean 103 mg vs. 529 mg, *p* ≤ 0.0001) and total BM cellularity (femur; mean 11.6 × 10^6^/mL vs. 15.7 × 10^6^/mL, *p* 0.0035) were also normalized, and while the extent of reticulin fibrosis was increased in *Jak2^RL^/Tet2^-/-^* mice compared to *Jak2^RL^*, mutant allele reversal resolved fibrosis in both mutational contexts **(Fig. 4g, Extended Data Fig. 8d-e)**. The reduction in mutant cell fraction, as was observed with single-mutant mice, persisted down to the level of HSPCs in tamoxifen treated *Jak2^RL^/Tet2^-/-^*mice, including within the LSK stem cell-enriched compartment (mean 28.7% vs. 73.7%, *p* ≤ 0.001) **(Fig. 4h, Extended Data Fig. 8f)**. In a subset of assayed *Jak2^RL^/Tet2^-/-^* mice following *Jak2^V617F^* deletion (4/9), we were unable to detect *Tet2^-/-^*knock-out bands in whole marrow at time of sacrifice, and cells harvested from *Jak2^RL^/Tet2^-/-^* recipient mice following oncogenic deletion were unable to serially replate indicating loss of self-renewal capacity in comparison to control double-mutant mice **(Fig. 4i, Extended Data Fig. 8g)**. These data support the notion that co-occurring loss-of-function mutations of *Tet2* do not dramatically alter reliance on JAK/STAT signaling for disease maintenance, and that despite the fitness advantage engendered by *Tet2* loss on MPN hematopoietic stem cells, the reductions in HSC fitness in the setting of *Jak2^V617F^* reversion suggest a unique dependency on oncogenic Jak2^V617F^ that renders double-mutant cells susceptible to eradication.

## Conclusion

Mutated kinases occur frequently in cancer and are amenable to targeted inhibition; however, mechanisms mediating acquired resistance have been observed for most targeted therapies.^38^ By contrast, current JAK inhibitors fail to eliminate JAK2^V617F^-mutant clones in MPN patients suggesting inadequate target inhibition and/or other genetic/non-genetic factors mediate JAK2^V617F^-mutant cell persistence in the setting of JAK inhibitor therapy.^5^ We show in preclinical models that there is an absolute requirement for Jak2^V617F^ in MPN cells and that mutant-specific targeting of Jak2^V617F^ abrogates MPN features, reduces mutant cell fraction, and extends overall survival with concomitant depletion of disease-sustaining stem cells within the HSPC compartment. Further, our data suggest that Jak2^V617F^ dependency persists even in the setting of antecedent mutations in epigenetic regulators, specifically *Tet2*. These data support the notion that improved targeting of JAK2 signaling and downstream effectors offers greater therapeutic potential than current JAK kinase inhibitors and that JAK2^V617F^ mutant-selective inhibition represents a potential curative strategy for the treatment of MPN patients. Moreover, we demonstrate the feasibility of our dual-recombinase system to evaluate oncogenic signaling dependencies *in vivo*, and we believe that a similar approach will allow us to assess oncogenic dependencies and mechanisms of mutant-mediated transformation across a spectrum of malignant contexts.

## METHODS

### Experimental animals

All animal studies were performed in accordance with institutional guidelines established by Memorial Sloan Kettering Cancer Center (MSKCC) under the Institutional Animal Care and Use Committee-approved animal protocol (#07-10-016) and the Guide for the Care and Use of Laboratory Animals (National Academy of Sciences 1996). All experimental animals were maintained on a 12 hour light-dark cycle with access to water and standard chow ad libitum. Veterinary staff provided regular monitoring and husbandry care. All mice had intact immune systems, were drug and test naïve, and had not been involved in previous procedures. Animals were monitored daily for signs of disease or morbidity, bleeding, failure to thrive, infection, or fatigue and sacrificed immediately if they exhibited any signs of the above. Mice harboring the *Jak2^RL^* allele were generated by Ingenious Targeting Laboratory (Ronkonoma, NY) in a C57BL/6 background. Specifically, a 8.86kb genomic DNA used to construct the targeting vector was first subcloned from a positively identified C57BL/6 BAC clone (RP23- 316C6). The region was designed such that the long homology arm (LA) extends ∼6 kb 5’ to the cluster of Lox2272-Rox-Rox12-Lox2272 sites, and the short homology arm (SA) extends about 2.2 kb 3’ to the Neo cassette and 3’ Rox12 site. The inversion cassette is in between the second set of Lox2272 and Rox sites, and it consists of the mutant exon 14* (V617F) and its flanking genomic sequences for correct splicing (SaE14*Sd). The inversion cassette replaces wild-type exon 14 and the same flanking genomic sequences included in the cassette. The BAC was sub-cloned into a ∼2.4kb pSP72 (Promega) backbone vector containing an ampicillin selection cassette for retransformation of the construct. Ten micrograms of the targeting vector was then linearized and transfected by electroporation of FLP C57Bl/6 (B6) embryonic stem cells. After selection with G418 antibiotic, surviving clones were expanded for PCR analysis to identify recombinant ES clones. After successful clone identification, the neomycin cassette was removed with a transient pulse of Cre recombinase and clones were reconfirmed following expansion. Finally, ES cells were injected in C57BL/6 mice via tetraploid complementation (NYU). *Tet2^f/f^* conditional knock-out mice, Cre-lox *Jak2^V617F^* knock-in mice, RC::RLTG reporter mice, Cre TdTomato reporter mice, and Ubc:CreER mice have been described previously.^6, 8, 21, 25, 39^ 6-8 week old female and male *Jak2^RL^* or *Jak2^RL^/Tet2^f/f^* donor mice were used for Dre electroporation knock-in experiments. Age-matched 6-10 week old female mice were used as donors for all transplant experiments (Ly5.1 Cd45.1 competitive or C57BL/6 non-competitive).

### Mouse genotyping

DNA was isolated using the DNeasy Blood & Tissue Kit (Qiagen, Germantown, MD). *Jak2^V617F^* knock-in genotyping was carried out with the following primers: FWD: 5’-GCCATCTTTCCAGCCTAAAATTAG-3’; REV: 5’-TCCAAAGAGTCTGTAAGTACAGAACT-3’ and with the following reaction conditions: 94°C for 3 minutes followed by 15 cycles of 94°C for 15s, 65°C for 15s, and 72°C for 30s decreasing by 1°C per cycle, and then followed by an additional 25 cycles of 94°C for 15s, 50°C for 15s, and 72°C for 30s. *Jak2^V617F^* knock-out genotyping was carried out using the following primers: FWD: 5’-GCCATCTTTCCAGCCTAAAATTAG-3’; REV: 5’-ACCAGTTGCTCCAGGGTTACACG-3’ and with the following reaction conditions: 94°C for 2 minutes followed by 30 cycles of 94°C for 30s, 53°C for 30s, and 72°C for 30s. Sequencing of the unrecombined Rox-lox locus was carried out using the following primers: FWD: 5’-AGGAGCATCGATGACTACATGATGAG- 3’; REV: 5’-AGACTCTCCACGGTCTCATCTACG-3’ and with the following reaction conditions: 98°C for 30 seconds followed by 35 cycles of 98°C for 10s, 65°C for 15s, and 72°C for 30s. *Tet2* genotyping were carried out using the following primers/conditions: FWD: 5’- AAGAATTGCTACAGGCCTGC-3’; REV: 5’-TTCTTTAGCCCTTGCTGAGC-3’; ExR: 5’-TAGAGGGAGGGGGCATAAGT-3’ and with the following reaction conditions: 94°C for 2 minutes followed by 39 cycles of 94°C for 35s, 58°C for 45s, and 72°C for 55s. Annotation of PCR genotyping results was carried out on a QIAxcel Advanced System (Qiagen) and analyzed using QIAxcel ScreenGel software (Qiagen). Sanger sequencing was performed by Genewiz (South Plainfield, NJ) and analyzed using Benchling software.

### Dre mRNA electroporation

Dre mRNA was purchased from TriLink Biotechnologies (San Diego, CA) and electroporation carried out using the Neon Transfection System (ThermoScientific) per the manufacturer’s protocol. Specifically, bone marrow donor cells were isolated from limb bones into phosphate buffered saline (PBS; pH 7.2) containing 2% fetal calf serum via centrifugation. After red blood cell (RBC) lysis, single-cell suspensions were depleted of lineage-committed hematopoietic cells using a Lineage Cell Depletion Kit according to manufacturer’s protocol (EasySep™, StemCell Technologies, Inc.). 2.5-3.0 × 10^6^ lineage-depleted bone marrow was then washed in PBS and then resuspended in 135 μL Buffer T to which 15 μL of Dre mRNA (at 1 μg/μL) was quickly added and electroporated at the following conditions: 1700V for 20ms x1 pulse. The cells were then pipetted into penicillin-streptomycin free StemSpan SFEM medium with thrombopoietin (TPO; 20 ng/mL; PeproTech) and stem cell factor (SCF; 20 ng/mL; PeproTech), cultured for two hours, and then subsequently harvested and washed/resuspended in PBS and transplanted via lateral tail vein injection into lethally irradiated (900cGy) 6-8 week old C57BL/6J recipient mice at approximately 4 × 10^5^ cells per recipient along with 50,000 un- electroporated wild-type whole bone marrow support cells. Double-mutant *Jak2^RL^/Tet2^f/f^* transplants/electroporations were carried out as above, except donor mice were dosed with tamoxifen (100 mg/kg by oral gavage daily x4; purchased from MedChemExpress) 6-8 weeks prior to harvest and excision confirmed prior to Dre electroporation.

### Transplantation assays and in vivo experiments

*Jak2^RL^* and *Jak2^RL^/Tet2^f/f^* lines were crossed to Ubc:CreER tamoxifen-inducible Cre lines and RLTG dual-recombinase reporter lines.^25, 39^ Primary recipient mice transplanted with Dre mRNA-recombined Ubc:CreER-*Jak2^RL^* or Ubc:CreER- *Jak2^RL^/Tet2^f/f^*bone marrow cells were bled every 3-4 weeks post-transplant to monitor disease status. Peripheral blood was isolated by submandibular bleeds and complete blood counts determined using a ProCyte Dx (IDEXX Laboratories, Westbrook, ME) per manufacturer’s instruction. For competitive repopulation assays, 1.2 × 10^6^ whole bone marrow from primary transplant recipient mice exhibiting MPN was harvested 6-8 weeks post-transplant and combined with age-matched 0.8 × 10^6^ Cd45.1 (Jackson Laboratories, Bar Harbor, ME) whole bone marrow and transplanted into 6-8 week old lethally irradiated Cd45.1 secondary recipient mice. Mice transplanted with Dre-recombined *Jak2^V617F^*cells demonstrating low Cd45.2 chimerism at baseline (<15%) and/or evidence of poor MPN cell engraftment were excluded from study cohorts. To induce Cre and delete *Jak2^V617F^*, mice were treated with tamoxifen (TAM; purchased from MedChemExpress) 100 mg/kg daily (dissolved in corn oil) by oral gavage x 4 followed by 14 days of TAM chow approx. 80 mg/kg daily (ENVIGO). Tamoxifen control studies were carried out using similar dosing schedules on 45.1 mice transplanted in competition with Dre-electroporated, Cre-negative *Jak2^RL^* MPN bone marrow cells. For terminal tissue isolation, mice were euthanized with CO2 asphyxiation, and tissues were dissected and fixed with 4% paraformaldehyde for histopathological analysis. For whole bone marrow isolation, the femurs, hips, and tibias were dissected and cleaned. Cells were then isolated using centrifugation at 8000xG for 1 minute followed by RBC lysis (BioLegend) for 10-15 minutes. Bone marrow cell numbers and viability were determined using an automated cell counter (ViCell Blu, Beckman Coulter). Spleen cell suspensions were generated by crushing whole spleen and filtering through a 70 μM filter. RBC lysis (BioLegend) was performed and cells were prepared for downstream processing or frozen.

### In vivo drug studies

For *in vivo* inhibitor studies, approximately 8 weeks following transplant, secondary transplant cohorts of lethally-irradiated mice transplanted with Ubc:CreER-*Jak2^RL^* bone marrow in competition with Cd45.1 marrow (as above) and exhibiting active MPN were bled and cohorted based on peripheral blood Cd45.2 chimerism and total WBC count to achieve congruency across treatment arms. Mice were then treated with ruxolitinib (60 mg/kg P.O. twice daily; dissolved in 20% Captisol in PBS; purchased from MedChemExpress), tamoxifen to delete *Jak2^V617F^* (as above; purchased from MedChemExpress), or vehicle. Investigators were not blinded to the identity of mice or samples. Mice were treated for a total of 6 weeks before timed sacrifice and marrow/spleen harvested as above.

### Bone marrow endothelial cell (BMEC) culture

Bone marrow cells were isolated from limb bones into FACS buffer (phosphate buffered saline [PBS] + 2% fetal bovine serum) via centrifugation. After RBC lysis, single-cell suspensions were depleted of lineage-committed hematopoietic cells using a Lineage Cell Depletion Kit according to manufacturer’s protocol (EasySep™, StemCell Technologies, Inc.). Subsequently, 50,000 of the resulting lineage^-^ cells were plated on a confluent monolayer of BMECs in a single well of a 12-well plate. Each well had 1 mL StemSpan SFEM (StemCell Technologies, Inc.) with 20 ng/mL recombinant murine SCF (PeproTech) in addition to the corresponding drug treatment: either 4-hydroxytamoxifen (4-OHT; Sigma Aldrich; stock concentration: 13 mM) or its vehicle, appropriately diluted in media to its final concentration (i.e., 0.01% (v/v) of ethanol (EtOH), or 200 nM, 400 nM or 1 μM of 4-OHT) (three replicates/condition). The BMECs were seeded two days before plating the lineage^-^ cells at a density of 100,000 cells/well. Co-cultures were maintained for a total of 7 days at 37°C and 5% CO2, with media being completely refreshed with the original SCF and drug/vehicle concentrations. 4-OHT or EtOH vehicle was added to the culture on day 1 and again on day 4. On day 7, total cells were harvested with Accutase (Biolegend) and cell numbers were determined via an automatic cell counter (ViCell Blu, Beckman Coulter). Cells were then stained with the desired antibody cocktail and phenotyped by flow cytometry.

### Flow cytometry and western blot

Following single cell preparation, murine peripheral blood, whole bone marrow, or spleen mononuclear cells were lysed for 10-15 minutes with RBC lysis buffer (BioLegend, San Diego, CA) and washed twice with FACS buffer. Cells were then resuspended in Fc (Cd16/32) block for 15 minutes and then subsequently stained with a cocktail comprised of antibodies targeting Cd3 (17A2), Cd45R/B220 (RA3-6B2), Gr-1 (RB6-8C5), Cd11b (M1/70), Cd45.2 (104), and Cd45.1 (A20) for 30 minutes. For hematopoietic stem/progenitor cell analysis, lysed bone marrow was stained with a cocktail of lineage markers along with antibodies against c-Kit (2B8), Sca-1 (D7), FcγRII/III (2.4G2), Cd34 (RAM34), Cd150 (9D1), and Cd48 (HM48-1). Erythroid progenitor flow was carried out on unlysed bone marrow or spleen with the addition of the following antibodies: Cd105, Cd71 (R17217), Cd41 (MWReg30), and Ter119 (Ter- 119). All FACS antibodies were purchased from BD, BioLegend, or eBioscience. Following antibody incubation, cells were washed with FACS buffer and resuspended in a DAPI-containing FACS buffer solution for analysis and sorting. Samples were run on a LSRFortessa (Becton Dickinson) using FACSDiva software and analyzed with FlowJo (Treestar, Ashland, OR, USA). For Western blot analysis, whole-cell protein extracts from harvested splenocytes were prepared using RIPA buffer (ThermoScientific, Rockford, IL) containing a protease/phosphatase inhibitor cocktail (Thermo Scientific). Protein quantification was performed using the Pierce BCA protein assay kit (ThermoScientific) and analyzed on a Cytation 3 plate reader (BioTek). Proteins were separated by NuPAGE 4-12% Bis-Tris Gel and transferred to a nitrocellulose membrane. The following antibodies were used: ≤-actin (Cell Signaling 4970S), STAT5 (Cell Signaling 94205S), and pSTAT5 (Cell Signaling, 9359S). Images were obtained using the ChemiDoc Imaging System (BioRad) and analyzed using ImageLab software (BioRad).

### Histology staining and immunohistochemistry (IHC), photography

Tibia and spleen samples were fixed in 4% paraformaldehyde for over 24 hours and then embedded in paraffin. Paraffin sections were cut on a rotary microtome (Mikrom International AG), mounted on microscope slides (ThermoScientific), and air-dried in an oven at 37°C overnight. After drying, tissue section slides were processed either automatically for hematoxylin and eosin (H&E) staining (COT20 stainer, Medite), or manually for reticulin staining. All samples and slide preparation, including immunohistochemistry was carried out at the Tri-Institutional Laboratory of Comparative Pathology (LCP) core facility. The following antibodies were used for immunohistochemistry: Mac1 (Cedarlane CL8941B, 1:100), Ter119 (BDBioscience, 550565 1:200), and phospho-44/42 MAPK (Erk1/2) (Cell Signaling 4376, 1:100). Pictures were taken at 100X, 200X and 400X (H&E, reticulin and respective IHC) magnification using an Olympus microscope and analyzed with Olympus Cellsens software. Tissue sections were formally evaluated by a hematopathologist (W. Xiao), including reticulin scoring.

### Assessment of apoptosis and viability

Apoptosis was measured by flow cytometry on a LSRFortessa (Becton Dickinson) cytometer with Annexin V PerCPCy5.5 antibody (BioLegend) in combination with the antibody cocktail (above) in Annexin binding buffer (BioLegend) at 1:50 dilution in combination with DAPI as live/dead cell stain.

### Colony forming assays

To assess colony formation and serial replating capacity, RBC-lysed 50,000 whole bone marrow cells were seeded in 1.5mL MethoCult M3434 (Stem Cell Technologies) with no additional supplemental cytokines in triplicate on 6 well plates and scored on day 8. For replating, cells were harvested and pooled and then re-seeded once more at 50,000 cells/well in 1.5mL MethoCult M3434 in triplicate.

### Serum cytokine profiling

Serum samples were diluted two-fold with PBS (pH 7.2) and stored at - 80°C until analysis. Cytokine levels were measured in duplicates (62.5μL each) by Eve Technologies Corporation (Mouse Cytokine Array/Chemokine Array 44-Plex, Calgary, AB, Canada).

### RNA sequencing (RNA-Seq) and data analysis

For gene expression analysis, secondary cohorts of lethally irradiated C57BL/6 mice transplanted with Ubc:CreER-*Jak2^RL^*-RLTG reporter bone marrow 8 weeks post-transplant and exhibiting MPN were treated with ruxolitinib (60mg/kg P.O. twice daily), tamoxifen (100mg/kg by oral gavage daily × 4 followed by 80mg/kg of TAM chow × 3 days) +/- vehicle (MPN control) for 7 days and then sacrificed. Lineage-depleted bone marrow was isolated and stained with an antibody cocktail containing a combination of lineage markers along with antibodies against c-Kit (2B8), Sca-1 (D7), FcγRII/III (2.4G2), and Cd34 (RAM34) for 30 minutes, washed, and then resuspended in FACS buffer containing DAPI as a live/dead stain. TdTomato+ (*Jak2^RL^* knock-in) or GFP+ (*Jak2^RL^* knock-out) LSKs and MEPs were then sorted on a FACSAria III directly into Trizol LS (Invitrogen) and stored at -80°C until processing. RNA was subsequently isolated using the Direct-Zol Microprep Kit (Zymo Research, R2061) according to manufacturer’s protocol and quantified using the Agilent High Sensitivity RNA ScreenTape (Agilent 5067-5579) on an Agilent 2200 TapeStation. cDNA was generated from 1 ng of input RNA using the SMART-Seq HT Kit (Takara 634455) at half reaction volume followed by Nextera XT (Illumina FC-131-1024) library preparation. cDNA and tagmented libraries were quantified using High Sensitivity D5000 ScreenTape (5067- 5592) and High Sensitivity D1000 ScreenTape respectively (5067- 5584). Libraries were sequenced on a NovaSeq at the Integrated Genomics Operation (IGO) at MSKCC. FASTQ files were mapped and transcript counts were enumerated using STAR (genome version mm10 and transcript version ensembl M13). Counts were input into R and RNA-sequencing analysis using DESeq2. Genes were filtered out prior to modeling in DESeq if they were not detected in all, with MEPs and LSKs modeled separately. Differentially expressed genes were identified with a log2-foldchange of 1 and an adjusted *p* value of 0.05. Gene set enrichment analysis was performed using the fgsea package at 100,000 permutations with genesets extracted from the msigdbr package. Single sample gene set enrichment analysis was performed using the gsva package. Figures were generated using ggplot2 and tidyheatmaps packages. Complete scripts can be found on github at https://github.com/bowmanr/goldilox.

### Mouse Assay for Transposase-Accessible Chromatin Sequencing (ATAC-Seq) and data analysis

Chromatin accessibility assays utilizing the bacterial Tn5 transposase were performed as described.^40^ Briefly, 5.0 × 10^4^ TdTomato+ (*Jak2^RL^* knock-in) or GFP+ (*Jak2^RL^* knock-out) cKit^+^ bone marrow cells from mice treated for 7 days with tamoxifen or an untreated MPN control cohort were sorted on a FACSAria III directly into PBS and subsequently lysed and incubated with transposition reaction mix containing PBS, Tagment DNA buffer, 1% Digitonin, 10% Tween-20, and Transposase (lllumina). Samples were then incubated for 30 minutes at 37°C in a thermomixer at 1000 rpm. Prior to amplification, samples were concentrated with the DNA Clean and Concentrator Kit-5 (Zymo). Samples were eluted in 20 μL of elution buffer and PCR-amplified using the NEBNext 2X Master Mix (NEB) for 11 cycles and sequenced on a NextSeq 500 (Illumina). All samples were processed at the Center for Epigenetics Research (CER) core facility at MSKCC. Libraries were sequenced on a NovaSeq at the Integrated Genomics Operation (IGO) at MSKCC. Data analysis was completed through in house scripts at the CER, in brief: reads were trimmed with ‘trim_galore’ and aligned to mouse genome mm9 using bowtie2 (default parameters). Duplicates were removed with the Picard tool ‘MarkDuplicates’, and peaks were called with MACS2, merged and used to create a full peak atlas. Read counts were tabulated over this atlas using featureCounts. Downstream differential enrichment testing was completed in DESeq2 with default normalization scheme. HOMER was used for known motif enrichment amongst the differentially enriched peaks as defined by a fold change of +/- 1.5 and an adjusted *p* value of 0.1.

### Human single-cell ATAC-Seq and data analysis

Single-cell ATAC-seq data was processed using cellranger-ATAC (v2.0.0) mkfastq. ATAC sequencing reads were then aligned to the hg38 reference genome using cellranger-ATAC count function. Fragment files generated by cellranger-ATAC were used as input for the ArchR^41^ (v1.0.0). For initial dimensionality reduction and patient data integration, the cell by genomic bin matrix was used as input for reciprocal latent semantic indexing (LSI) as calculated by the Signac (v1.1.1). Transcription factor motif accessibility z-scores were calculated with ChromVAR^42^ (v1.8.0). The earliest HSPCs (cluster HSPC1, see Myers, R. and Izzo, F. *et al*., bioRxiv, 2022) were subset for downstream analysis, and statistical comparisons of motif accessibility for NFKB1, REL, FOS, and JUN transcription factors were performed via linear mixture model including patient identity as random effect to account for potential technical confounders arising from sample-specific batch effects. For heatmap representation, motif accessibility z-scores were used as input and the pheatmap (v1.0.12) R package was used.

### Quantitative real-time PCR

Total RNA was extracted from magnetic-bead isolated cKit^+^ bone marrow (Miltenyi Biotec) using the Direct-zol RNA extraction kit (Zymo) per manufacturers’ protocols respectively. Complementary DNA was then reverse transcribed using the Verso cDNA Synthesis kit (ThermoFisher Scientific). *Ybx1* expression was evaluated by quantitative reverse-transcription (qRT) PCR using Taqman probes purchased from ThermoFisher (Mm00850878_g1) on the RealPlex thermocycler (ThermoFisher Scientific, Fairlawn, NJ).

### Statistical analysis

Statistical analyses were performed using Student’s *t-*test (normal distribution) using GraphPad Prism version 6.0h (GraphPad Software, San Diego, CA) unless otherwise noted. Kaplan-Meier curves were determined using the log-rank test. *P*<0.05 was considered statistically significant. The number of animals, cells and experimental replication can be found in the respective figure legends.

### Data availability

All raw and processed sequencing data is made available at https://github.com/bowmanr/goldilox and via the NCBI Gene-Expression Omnibus (GEO).

## ACKNOWLEDGEMENTS

We are grateful to members of the Levine Lab for their discussion of the work. We would also like to acknowledge Dr. Alex Joyner (MSKCC), Dr. Patricia Jensen (NIH) and Dr. Tudor Badea (NIH) for discussion on Dre-Rox technology, as well as Sara Meyer and Sime Brkic (University Hospital Basel Switzerland) for their technical advice and support. This work was supported by the National Cancer Institute award P01 CA108671 (R.L.L.). R.L.L. was supported by a Leukemia and Lymphoma Society Scholar award. A.J.D. is a William Raveis Charitable Fund Physician-Scientist of the Damon Runyon Cancer Research Foundation (PST-24-19). He also has received funding from the National Institute of Health (T32CA009207), American Association of Cancer Research (17-40-11-DUNB) and the American Association of Clinical Oncology. R.L.B. was supported by a Damon Runyon-Sohn Fellowship and the National Cancer Institute (K99CA248460). R.M.M. is supported by a Medical Scientist Training Program grant from the National Institute of General Medical Sciences of the National Institutes of Health under award number T32GM007739 to the Weill Cornell/Rockefeller/Sloan Kettering Tri-Institutional MD-PhD Program and by the Weill Cornell Medicine NYSTEM Training Program under award number C32558GG. F.I. is supported by the American Society of Hematology Fellow-to-Faculty Scholar Award. D.A.L. is supported by the Burroughs Wellcome Fund Career Award for Medical Scientists, Valle Scholar Award, Leukemia Lymphoma Scholar Award, Mark Foundation Emerging Leader Award and the National Institutes of Health Director’s New Innovator Award (DP2-CA239065). This work was also supported by the Tri-Institutional Stem Cell Initiative, the National Heart Lung and Blood Institute (R01HL145283; R01HL157387-01A1) and the National Human Genome Research Institute, Center of Excellence in Genomic Science (RM1HG011014). Studies supported by MSK core facilities were supported in part by MSKCC Support Grant/Core Grant P30 CA008748 and the Marie-Josée and Henry R. Kravis Center for Molecular Oncology. R.L.L. is also supported by a Leukemia & Lymphoma Society Specialized Center of Research grant.

## AUTHOR CONTRIBUTIONS

A.D., R.L.B. and R.L.L. conceived the project, designed the experiments, and analyzed the data. In vivo work was performed primarily by A.D. with technical assistance from Y.P., A.K., W.J.K., A.N, M.B, M.F, S.C., L.C., B.W., W.A., S.M., S.E., T.M.. Additional project design provided by A.V.. Hematopathology was formally interpreted by W.X.. Bone marrow endothelial experiments were performed by A.D., M.W., and I.F.M.. R.L.B., J.Y, and R.K. performed the computational analysis. Single-cell human ATAC data was performed and analyzed primarily by F.I., R.M.M, with support from D.L.. A.D., R.L.B., and R.L.L. contributed to the initial manuscript drafts. All authors reviewed and commented on the final manuscript. Funding acquisition: R.L.L., D.L., A.D., and R.L.B..

## MATERIALS AND CORRESPONDENCE

Correspondence and requests for materials should be addressed to R.L.L.

---

**Extended Data Figure 1:**
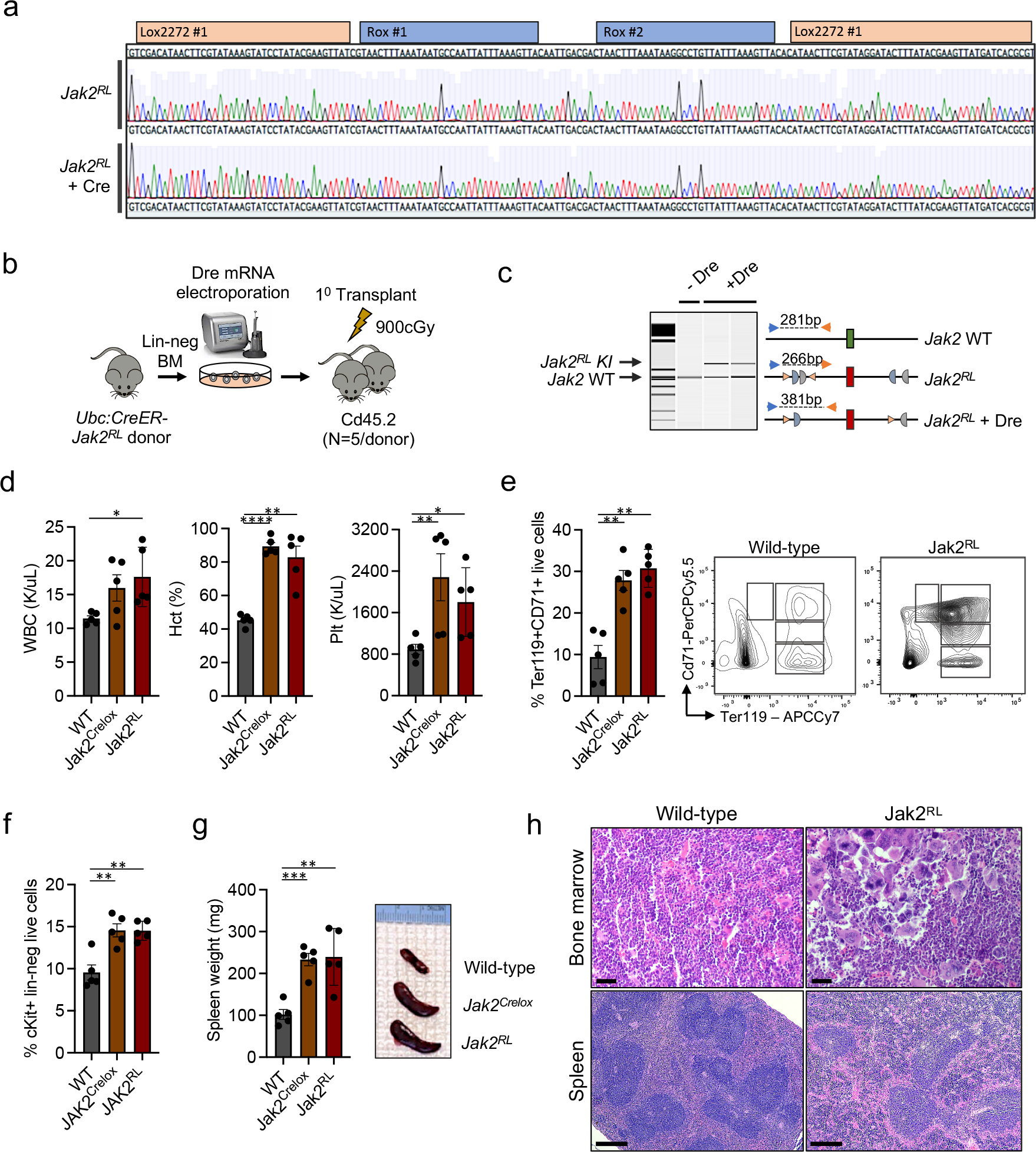
Phenotypic characterization and validation of *Jak2^RL^* activation *in vivo.* **a.** Sanger sequencing of the *Jak2^RL^* locus from genomic DNA of sorted Cre TdTomato+ reporter bone marrow cells following tamoxifen administration (*Jak2^RL^*+ Cre) compared to the non-recombined *Jak2^RL^* locus (*Jak2^RL^*) demonstrating retainment of *lox*/*rox* sites despite prior Cre exposure. Representative of *n* = 3 individual biological replicates. **b.** Schematic of the Dre mRNA *Jak2^RL^* bone marrow electroporation transplant protocol. Lin-neg BM: lineage-negative bone marrow; cGy: centigray. **c.** Representative polymerase chain reaction (PCR) of genomic DNA isolated from peripheral blood mononuclear cells in primary transplant recipients following Dre mRNA electroporation (+Dre) confirming the presence of *Jak2^V617F^* knock-in bands (KI) in relation to the *Jak2* wild-type (WT) allele in comparison to unrecombined (-Dre) *Jak2^RL^* cells. **d.** Peripheral blood counts of wild-type (WT) vs. *Jak2^RL^*knock-in mice in comparison to the previously published Cre-lox conditional *Jak2^V617F^* knock-in mouse model (*Jak2^Crelox^*)^8^ at timed sacrifice of 16 weeks: white blood cells (WBC; left panel), hematocrit (Hct; middle panel), platelets (Plt; right panel) (*n* = 5 per arm; mean ± s.d.). \**p* ≤ 0.05, \*\**p* ≤ 0.01, \*\*\*\**p* ≤ 0.0001. **e.** Ter119^+^Cd71^+^ erythroid precursor fractions (left panel) and representative flow cytometry plots (right panel) from harvested spleen cells of *Jak2^RL^* knock-in mice in comparison to wild-type (WT) and the *Jak2^Crelox^* knock-in model (*n* = 5 per arm; mean ± s.e.m). Black boxes on flow plots denote Ter119^+^ erythroid progenitor stages I-IV. \*\**p* ≤ 0.01. **f.** Bone marrow myeloid progenitor (Lineage^-^ cKit^+^Sca1^-^) frequencies of *Jak2^RL^* knock-in mice in comparison to wild-type (WT) and the *Jak2^Crelox^* knock-in model (*n* = 5 per arm; mean ± s.e.m). \*\**p* ≤ 0.01. **g.** Spleen weights (left panel) and representative spleens (right panel) of *Jak2^RL^* knock-in mice in comparison to wild-type (WT) and the *Jak2^Crelox^* knock-in model at timed sacrifice 16 weeks post-transplant (*n* = 5 per arm; mean ± s.d.). \*\**p* ≤ 0.01, \*\*\**p* ≤ 0.001. **h.** Hematoxylin and eosin (H&E) stains of bone marrow and spleen of *Jak2^RL^* knock-in mice in comparison to wild-type controls at timed sacrifice 16 weeks post-transplant. Representative micrographs of *n* = 5 individual mouse replicates per arm. Bone marrow images represented at 400X magnification (scale bar 20μm); spleen images 100X (scale bar 100μm). **d-h.** Representative of an *n* = 3 independent transplants.

**Extended Data Figure 2:**
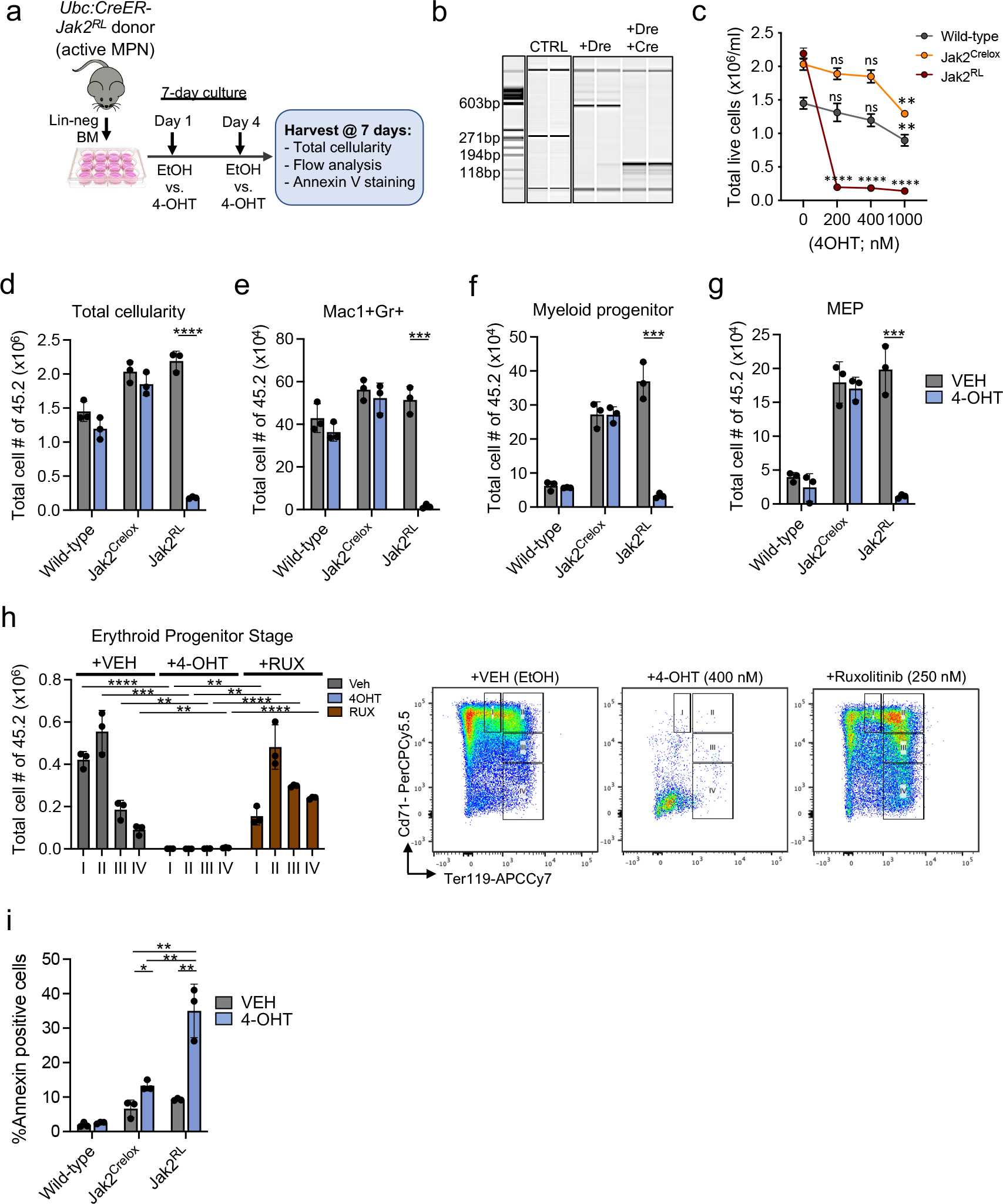
Functional consequences of *Jak2^V617F^* reversion *ex vivo.* **a.** Schematic of the bone marrow endothelial cell (BMEC) co-culture assay.^16^ Lin-neg BM: lineage-negative BM; EtOH: ethanol vehicle; 4-OHT: 4-hydroxy-tamoxifen. **b.** Representative *Jak2^V617F^* knock-in and knock-out genotyping by polymerase chain reaction (PCR) of genomic DNA isolated from cultured bone marrow cells following Dre-mediated knock-in (+Dre) and/or subsequent Cre-mediated deletion (+Dre+Cre) in comparison to the unrecombined *Jak2^RL^*allele (CTRL). Representative bands of *n* = 4 individual replicates per arm. **c.** Total cell output of cultured wild-type (gray) vs. *Jak2^RL^* knock-in (maroon) cells in comparison to Cre-inducible *Jak2^V617F^* knock-in (*Jak2^Crelox^*; gold)^8^ cells following 7 days of culture over BMECs in the presence of increasing doses (0 nM - 1000 nM) of 4-hydroxy-tamoxifen (4-OHT) (*n* = 3 technical replicates each; mean ± s.d.). ns: not significant, \*\**p* ≤ 0.01, \*\*\*\**p* ≤ 0.0001. **d.** Total cell, **e.** Mac1^+^Gr1^high^ mature neutrophil, **f.** Myeloid progenitor (Lineage^-^cKit^+^Sca1^-^), and **g.** Megakaryocytic-erythroid progenitor (MEP; Lineage^-^cKit^+^Sca1^-^Cd34^-^Fcg^-^) cell output of *Jak2^RL^* knock-in cells in comparison to wild-type vs. Cre-inducible *Jak2^Crelox^*MPN cells following 7 days of culture over BMECs in the presence of vehicle (VEH) vs. 400 nM 4-hydroxy-tamoxifen (4-OHT) (*n* = 3 technical replicates each; mean ± s.d.). \*\*\**p* ≤ 0.001, \*\*\*\**p* ≤ 0.0001. **h.** Total erythroid precursor cell numbers by Ter119/Cd71 erythroid progenitor flow cytometry (I-IV; left panel) and representative Ter119/Cd71 flow plots (right panel) of cultured *Jak2^RL^* MPN cells following 7 days of culture over BMECs in the presence of vehicle (VEH) vs. 400 nM 4-hydroxy-tamoxifen (4-OHT) vs. 250 nM ruxolitinib (RUX) (*n* = 3 technical replicates each; mean ± s.d.). \*\**p* ≤ 0.01, \*\*\**p* ≤ 0.001, \*\*\*\**p* ≤ 0.0001. **i.** Percentages of Mac1^+^Gr1^high^ Annexin V+ apoptotic cells of *Jak2^RL^* cells in comparison to wild-type vs. *Jak2^Crelox^* cells following 7 days of culture over BMECs in the presence of vehicle (VEH) vs. 200 nM 4-hydroxy-tamoxifen (4-OHT) (*n* = 3 technical replicates each; mean ± s.d.). \**p* ≤ 0.05, \*\**p* ≤ 0.01. **c-i.** Representative of *n* = 2 independent experiments.

**Extended Data Figure 3:**
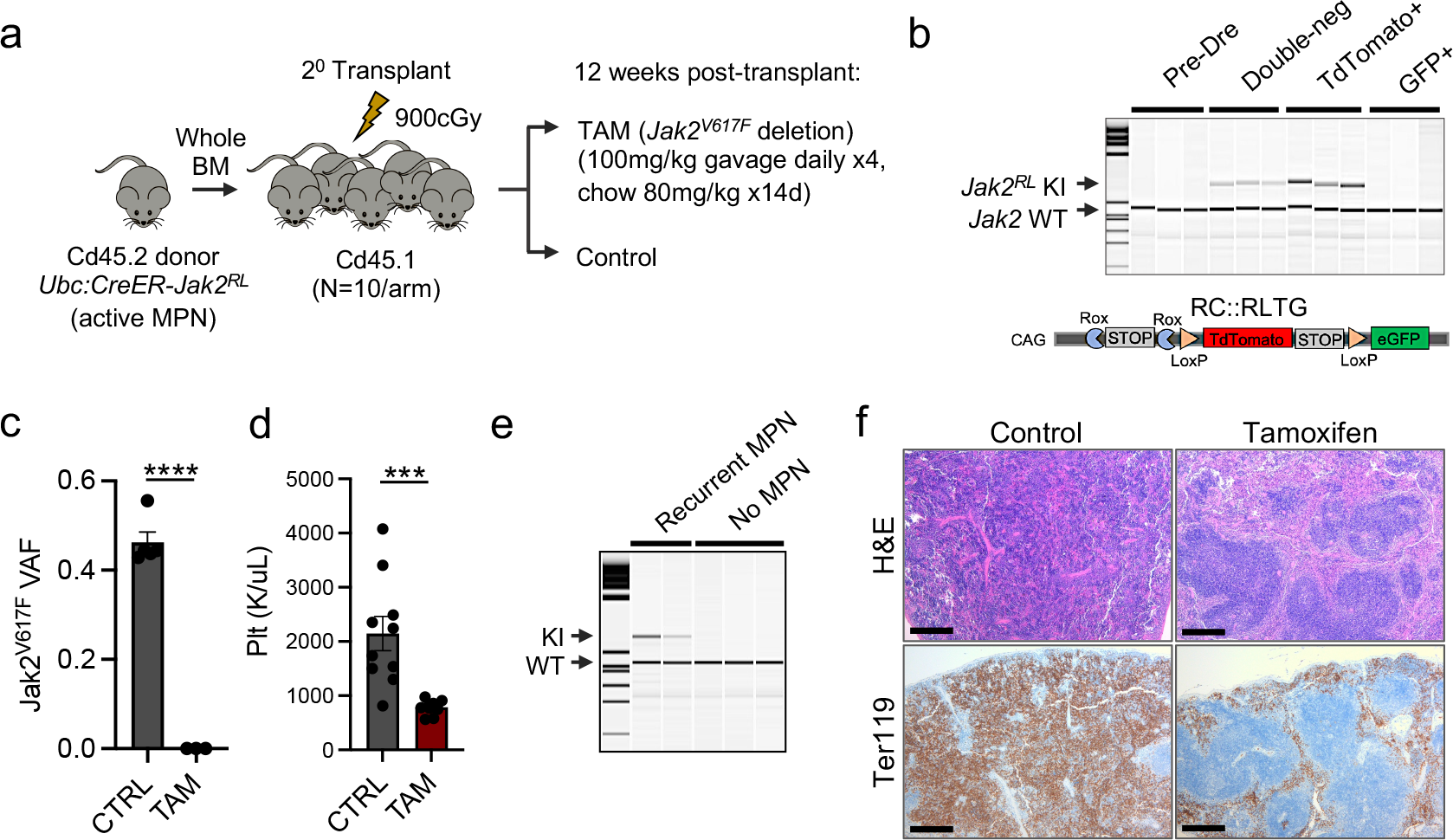
Functional characterization of *Jak2^RL^* knock-in/knock-out *in vivo.* **a.** Schematic of the experimental set up for the *Jak2^RL^* knock-in/knock-out transplants. BM: bone marrow; TAM: tamoxifen; cGy: centigray. b. Representative polymerase chain reaction (PCR) demonstrating presence or loss of *Jak2^RL^* knock-in (KI) bands in relation to the *Jak2* wild-type (WT) allele from reporter-sorted whole bone marrow mononuclear cells based on different recombined states: Pre-Dre: unrecombined *Jak2^RL^* cells; double-neg: reporter-negative cells following Dre recombination; TdTomato+: *Jak2^V617F^* knock-in population following Dre recombination; GFP+: green fluorescent protein (GFP) population: *Jak2^V617F^*-deleted population (i.e. Dre followed by Cre). c. *Jak2^V617F^* mutant RNA transcript variant allele frequency (VAF) from isolated megakaryocytic-erythroid progenitor (MEP; Lineage^-^cKit^+^Sca1^-^Cd34^-^Fcg^-^) cells 7 days following tamoxifen (TAM; *Jak2^V617F^* deletion) treatment in comparison to MPN (CTRL) mice (*n* = 3-5 each; mean ± s.d.). \*\*\*\**p* ≤ 0.0001. d. Platelet (Plt) counts of MPN (CTRL) vs. tamoxifen (TAM; *Jak2^V617F^*-deleted) treated mice at timed sacrifice 24 weeks (*n* ≥ 10 each; mean ± s.e.m). Representative of *n* = 2 independent transplants. \*\*\**p* ≤ 0.001. e. Representative polymerase chain reaction (PCR) demonstrating presence of *Jak2^V617F^* knock-in (KI) bands in relation to the *Jak2* wild-type (WT) allele from isolated bone marrow mononuclear cells of transplant recipients previously treated with tamoxifen (*Jak2*^V617F^ deletion) and exhibiting phenotypic features of recurrent MPN (*n* = 2) in comparison to those with no recurrent MPN (representative of *n* = 8 biological replicates). f. Representative hematoxylin and eosin (H&E) stains and Ter119 immunohistochemistry of sectioned spleen of MPN control vs. tamoxifen (*Jak2*^V617F^-deleted) treated mice at timed sacrifice 24 weeks. Representative micrographs of *n* = 6 individual mouse replicates per arm. All images represented at 100X magnification. Scale bar: 100μm.

**Extended Data Figure 4:**
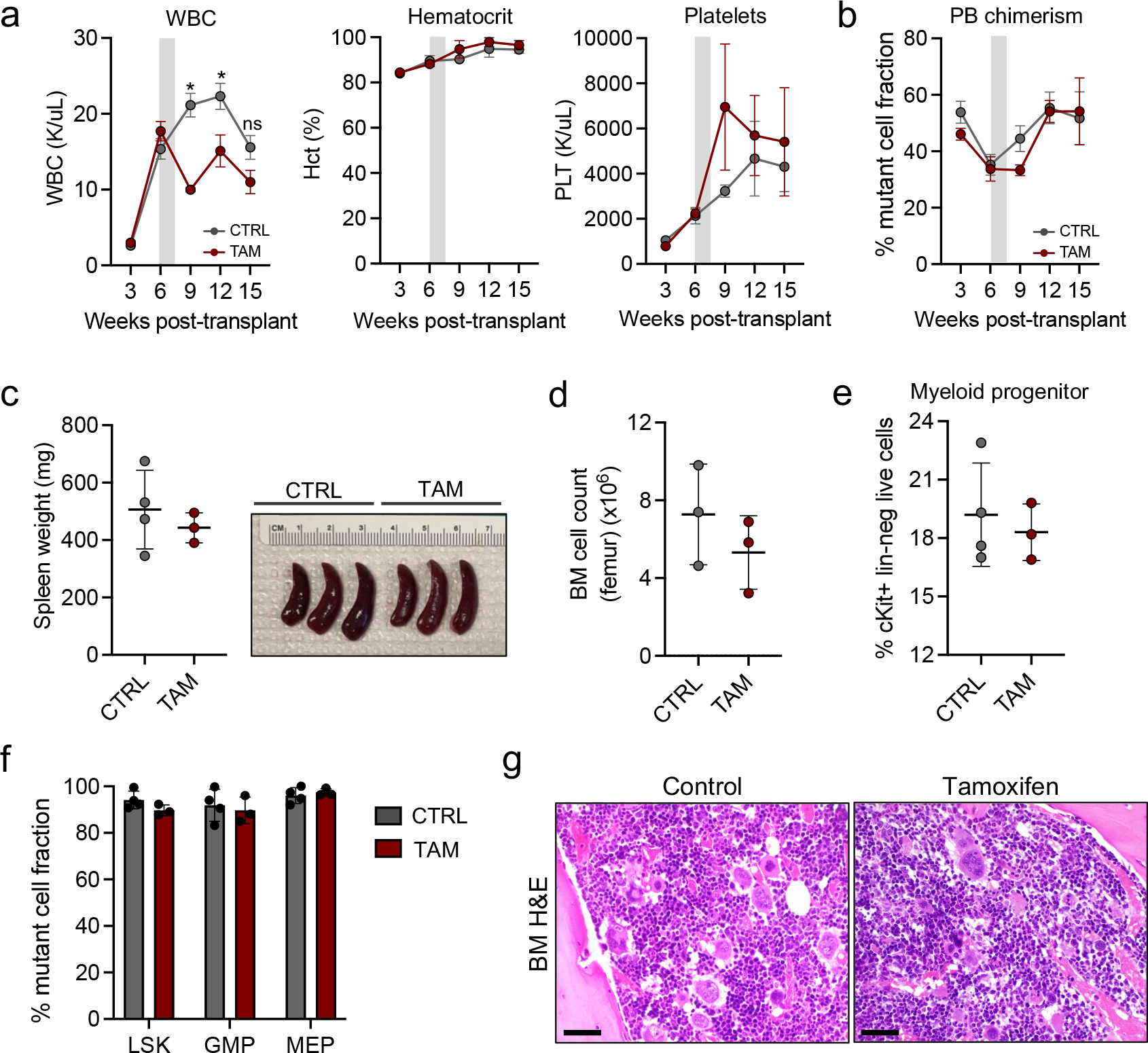
Assessment of tamoxifen toxicity on *Jak2^RL^*knock-in cells. **a.** Peripheral blood count trend (weeks 0-15) of MPN (CTRL) vs. tamoxifen (TAM) treated mice transplanted with Cre-negative *Jak2^RL^* knock-in cells in competition with Cd45.1 support: white blood cells (WBC; left panel), hematocrit (Hct; middle panel), platelets (PLT; right panel) (*n* = 3- 4 each; mean ± s.e.m). Gray bars represent duration of TAM pulse/chow administration. ns = not significant, \**p* ≤ 0.05. **b.** Peripheral blood (PB) mutant Cd45.2 percent chimerism trend (weeks 0- 15) of MPN (CTRL) vs. tamoxifen (TAM) treated mice transplanted with Cre-negative *Jak2^RL^* knock-in cells in competition with Cd45.1 support (*n* = 3-4 each; mean ± s.e.m). Gray bar represents duration of TAM pulse/chow administration. **c.** Spleen weights (left panel) and representative isolated spleens (right panel) of MPN (CTRL) vs. tamoxifen (TAM) treated mice transplanted with Cre-negative *Jak2^RL^* knock-in cells in competition with Cd45.1 support at timed sacrifice 15 weeks (*n* = 3-4 each; mean ± s.e.m). **d.** Total bone marrow (BM) cellularity (femur) and **e.** Myeloid progenitor (Lineage^-^cKit^+^Sca1^-^) frequencies of MPN (CTRL) vs. tamoxifen (TAM) treated mice transplanted with Cre-negative *Jak2^RL^* knock-in cells in competition with Cd45.1 support at timed sacrifice 15 weeks (*n* = 3-4 each; mean ± s.e.m). **f.** Bone marrow mutant cell fraction within LSK (Lineage^-^Sca1^+^cKit^+^), granulocytic-monocytic progenitor (GMP; Lineage^-^cKit^+^Sca1^-^Cd34^+^Fcg^+^), and megakaryocytic-erythroid progenitor (MEP; Lineage^-^ cKit^+^Sca1^-^Cd34^-^Fcg^-^) compartments of MPN (CTRL) vs. tamoxifen (TAM) treated mice transplanted with Cre-negative *Jak2^RL^* knock-in cells in competition with Cd45.1 support at timed sacrifice 15 weeks (*n* = 3-4 each; mean ± s.e.m.). **g.** Hematoxylin and eosin (H&E) stains of bone marrow (BM) of MPN (CTRL) vs. tamoxifen (TAM) mice transplanted with Cre-negative *Jak2^RL^* knock-in cells in competition with Cd45.1 support at timed sacrifice 15 weeks. Representative micrographs of *n* = 3-4 individual mouse replicates. Images represented at 400X magnification. Scale bar: 20μm. **a-g.** Representative of *n* = 3 independent experiments.

**Extended Data Figure 5:**
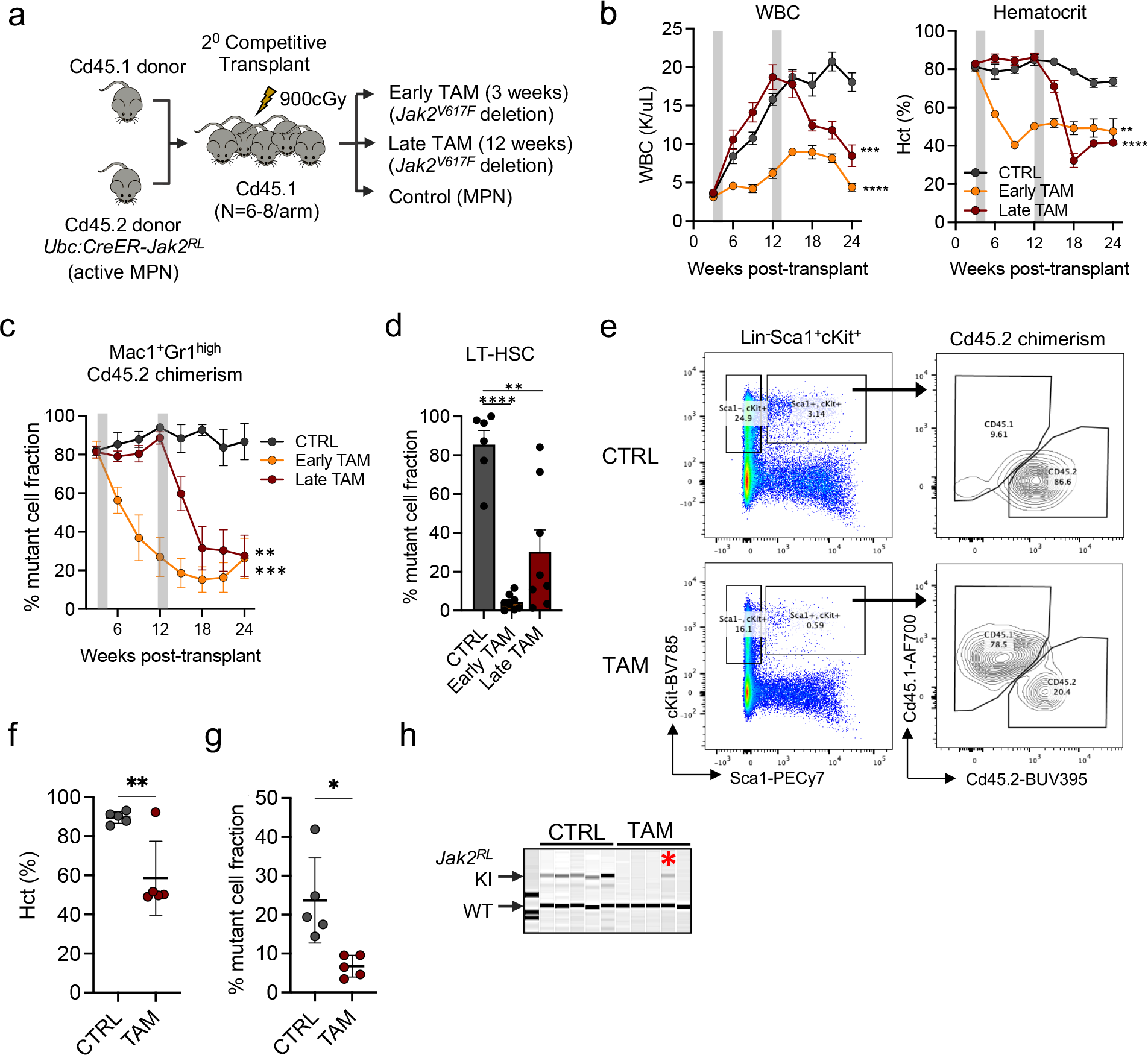
Effects of *Jak2^V617F^* deletion on MPN stem cell fitness and disease transplantability. **a.** Schematic of the experimental set up for the *Jak2^RL^* competitive transplant studies. TAM: tamoxifen; cGy: centigray. **b.** Peripheral blood count trends (weeks 0-24) of early (3 weeks post-transplant) tamoxifen (TAM; *Jak2^V617F^*-deleted) treated (gold) and late (12 weeks post-transplant) tamoxifen treated (maroon) mice in comparison to MPN (CTRL; gray) mice: white blood cells (WBC; left panel), hematocrit (Hct; right panel) (*n* = 6-8 per arm; mean ± s.e.m). Gray bars represent duration of TAM pulse/chow administration. Representative of *n* = 2 independent transplants. \*\**p* ≤ 0.01, \*\*\**p* ≤ 0.001, \*\*\*\**p* ≤ 0.0001. **c.** Peripheral blood Mac1^+^Gr1^high^ mutant Cd45.2 percent chimerism trend (weeks 0-24) of early (3 weeks post-transplant) tamoxifen (TAM; *Jak2^V617F^*-deleted) treated (gold) and late (12 weeks post-transplant) tamoxifen treated (maroon) mice in comparison to MPN (CTRL; gray) mice (*n* = 6-8 each, mean ± s.e.m.). Gray bars represent duration of TAM pulse/chow administration. Representative of *n* = 2 independent transplants. \*\**p* ≤ 0.01, \*\*\**p* ≤ 0.001. **d.** Bone marrow mutant Cd45.2 percentage within the long-term hematopoietic stem cell (LT-HSC; Lineage^-^ Sca1^+^cKit^+^Cd150^+^Cd48^-^) compartment of early (3 weeks post-transplant) tamoxifen (TAM; *Jak2^V617F^*-deleted) treated and late (12 weeks post-transplant) tamoxifen treated mice in comparison to MPN (CTRL) mice at timed sacrifice 24 weeks (*n* = 6-8 each; mean ± s.e.m.). Representative of *n* = 2 independent transplants. \*\**p* ≤ 0.01, \*\*\*\**p* ≤ 0.0001. **e.** Representative flow cytometry plots demonstrating mutant Cd45.2 to competitor Cd45.1 percentage of gated LSK (Lineage^-^Sca1^+^cKit^+^) cells of MPN (CTRL) vs. tamoxifen (TAM; *Jak2^V617F^*-deleted) treated mice at timed sacrifice 24 weeks. Representative of *n* = 6-8 biological replicates per arm. **f.** Hematocrit (Hct) and **g.** Peripheral blood mutant Cd45.2 fraction of recipient mice transplanted with MPN (CTRL) vs. tamoxifen (TAM; *Jak2^V617F^*-deleted) treated unfractionated donor bone marrow cells at 12 weeks post-transplant (*n* = 5 each; mean ± s.e.m). Representative of *n* = 2 independent experiments. \**p* ≤ 0.05, \*\**p* ≤ 0.01. **h.** Polymerase chain reaction (PCR) demonstrating presence of a *Jak2^V617F^* knock-in (KI) band in relation to the *Jak2* wild-type (WT) allele in 1 of 5 recipients (red asterisk) transplanted with tamoxifen (TAM; *Jak2^V617F^*-deleted) treated donor cells demonstrating recurrence of MPN phenotype in comparison to donor mice transplanted with MPN (CTRL) bone marrow donor cells.

**Extended Data Figure 6:**
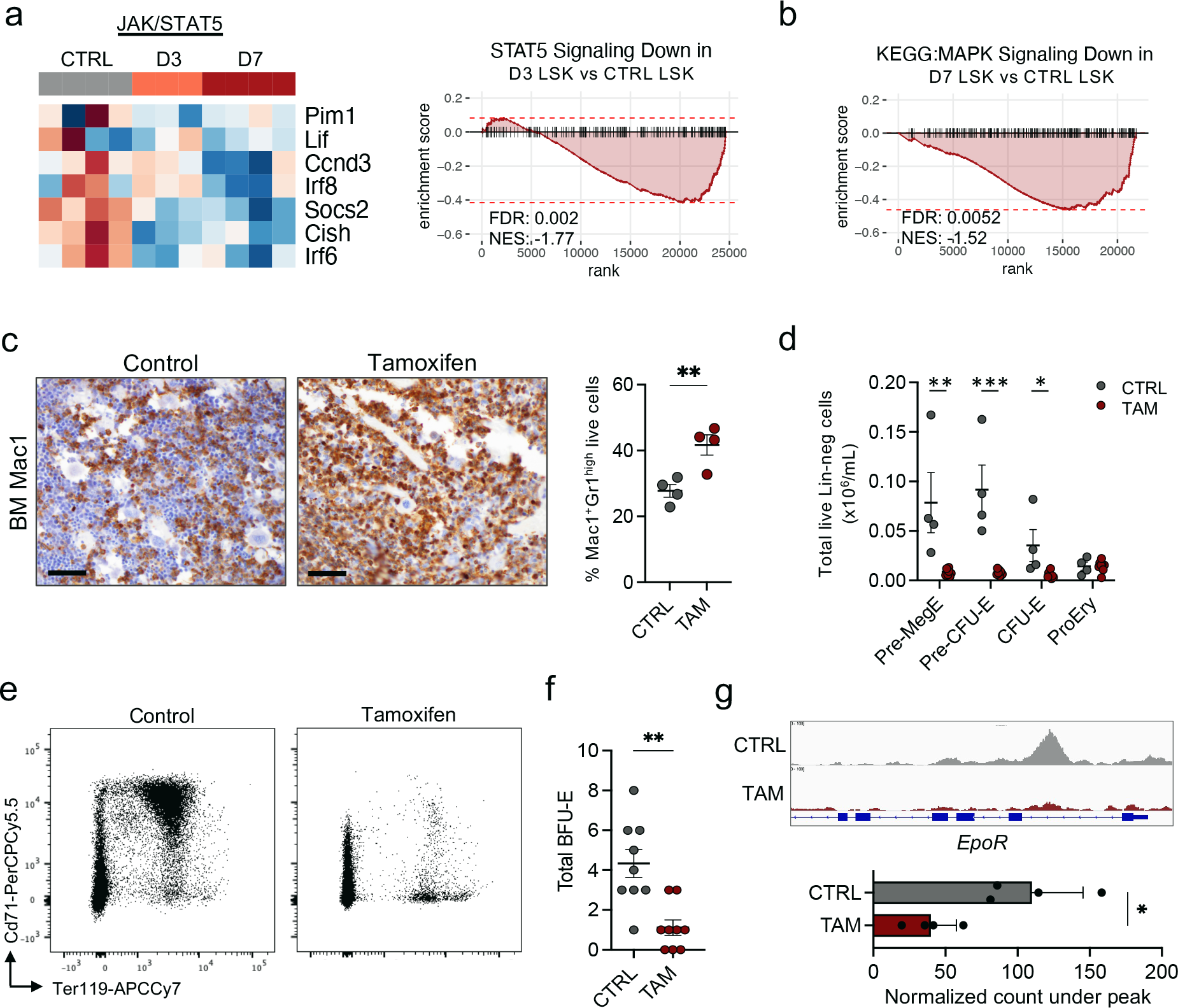
Acute phenotypic and transcriptional changes following *Jak2^V617F^* reversion. **a.** Heatmap (left panel) and gene set enrichment analysis (GSEA; right panel) demonstrating negative enrichment of STAT5 gene targets, including negative regulators of JAK/STAT signaling, in LSK (Lineage^-^Sca1^+^cKit^+^) cells harvested 3 days (D3) +/- 7 days (D7) following initiation of tamoxifen (*Jak2^V617F^*deletion) treatment in comparison to MPN (CTRL) cells. **b.** GSEA demonstrating negative enrichment in KEGG:MAPK signaling targets from harvested LSKs 7 days following initiation of tamoxifen treatment in comparison to MPN control (CTRL) LSKs. **c.** Mac1^+^ immunohistochemistry (IHC) of sectioned bone marrow (BM; left panel) and bone marrow Mac1^+^Gr1^high^ frequencies by flow cytometry (right panel) of MPN (CTRL) vs. tamoxifen (TAM; *Jak2^V617F^*-deleted) treated mice at 7 days following treatment (*n* = 4 each; mean ± s.e.m.). Representative micrographs of *n* = 4 individual mouse replicates each. Images represented at 400X magnification. Scale bar: 20μm. \*\**p* ≤ 0.01. **d.** Total erythroid progenitor bone marrow cell numbers, and **e.** Representative flow cytometry plots of Ter119/Cd71 erythroid progenitor populations of MPN (CTRL) vs. tamoxifen (TAM; *Jak2^V617F^*-deleted) treated mice 7 days following treatment (*n* = 4-7 per arm; mean ± s.e.m.). \**p* ≤ 0.05, \*\**p* ≤ 0.01, \*\*\**p* ≤ 0.001. **f.** Burst-forming unit erythroid (BFU-E) colony output of MPN (CTRL) vs. tamoxifen (TAM; *Jak2^V617F^*-deleted) treated marrow isolated 7 days following TAM administration and scored at day 8 after plating (each sample plated in triplicate, mean ± s.e.m.). **g.** Normalized ATAC-Seq signal at the *EpoR* locus (top panel) with associated quantified normalized peak counts (bottom panel) demonstrating decreased accessibility in tamoxifen (TAM; *Jak2^V617F^*-deleted) treated cKit^+^ bone marrow cells 7 days following initiation of TAM in comparison to MPN (CTRL) cells. **d-f.** Representative of *n* = 2 independent experiments.

**Extended Data Fig. 7:**
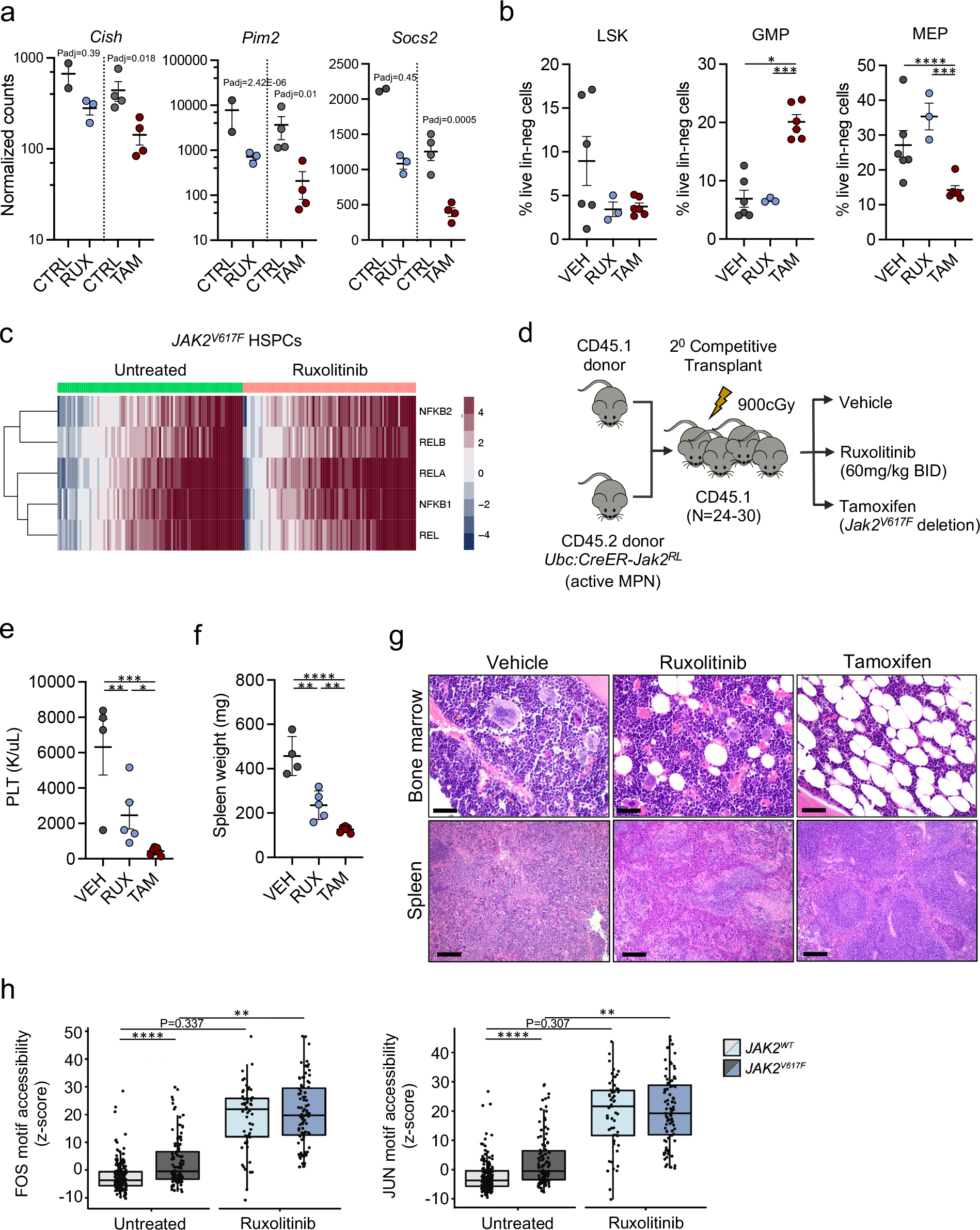
Differential responses of *Jak2^V617F^* deletion compared to JAK inhibitor therapy *in vivo.* **a.** Normalized read counts of representative JAK/STAT targets from LSK (Lineage^-^Sca1^+^cKit^+^) cells isolated 7 days following ruxolitinib (RUX) vs. tamoxifen (TAM; *Jak2^V617F^* deletion) treatment compared to MPN controls (CTRL) (*n* = 2-4 per arm; mean ± s.e.m.). LSK (Lineage^-^Sca1^+^cKit^+^; left panel), granulocytic-monocytic progenitor (GMP; Lineage^-^ cKit^+^Sca1^-^Cd34^+^Fcg^+^; middle panel), and megakaryocytic-erythroid progenitor (MEP; Lineage^-^ cKit^+^Sca1^-^Cd34^-^Fcg^-^; right panel) bone marrow frequencies of vehicle (VEH), ruxolitinib (RUX), or tamoxifen (TAM; *Jak2^V617F^*-deleted) treated mice 7 days following respective treatment (*n =* 3- 5 per arm; mean ± s.e.m.). Data representative of *n* = 2 independent experiments. \**p* ≤ 0.05, \*\*\**p* ≤ 0.001, \*\*\*\**p* ≤ 0.0001. **c.** Heatmap of single-cell ATAC-seq motif accessibilities for NFκB transcription factors for untreated (*n* = 105 cells from 4 patients) and ruxolitinib treated (*n* = 87 cells from 3 patients) human *JAK2^V617F^*mutant hematopoietic stem/progenitor cells (HSPCs) (from Myers, R. and Izzo, F. *et al*., bioRxiv, 2022). **d.** Schematic of the competitive transplant set up for the *Jak2^RL^ in vivo* drug studies. cGy: centigray. **e.** Platelet (PLT) counts and **f.** Spleen weights of vehicle (VEH), ruxolitinib (RUX), or tamoxifen (TAM; *Jak2^V617F^*-deleted) treated mice at the conclusion of the 6-week *in vivo* trial (*n* ≥ 4 each; mean ± s.e.m). \**p* ≤ 0.05, \*\**p* ≤ 0.01, \*\*\**p* ≤ 0.001, \*\*\*\**p* ≤ 0.0001. Representative of *n* = 3 independent experiments. **g.** Hematoxylin and eosin (H&E) stains of bone marrow and spleen sections from vehicle, ruxolitinib, or tamoxifen (*Jak2^V617F^*-deleted) treated mice at the conclusion of the 6-week *in vivo* trial. Representative micrographs of *n* = 4 individual mouse replicates per arm. Bone marrow images represented at 400X magnification (scale bar 20μm); spleen images 100X (scale bar 100μm). **h.** Box plots of single-cell ATAC-seq motif accessibility for FOS (left panel) and JUN (right panel) transcription factors for untreated (*n* = 188 cells from 4 patients; light gray) or ruxolitinib treated (*n* = 55 cells from 3 patients; light blue) *JAK2* wild-type human HSPCs in comparison to untreated (*n* = 105 cells from 4 patients; dark gray) or ruxolitinib-treated (*n* = 87 cells from 3 patients; dark blue) *JAK2^V617F^*-mutant HSPCs (from Myers, R.M. and Izzo, F. *et al*., bioRxiv, 2022). *P* values indicated are from linear mixture model explicitly modeling patient identity as random effect to account for patient-specific effects, followed by likelihood ratio test. \*\**p* ≤ 0.01, \*\*\*\**p* ≤ 0.0001.

**Extended Data Fig. 8:**
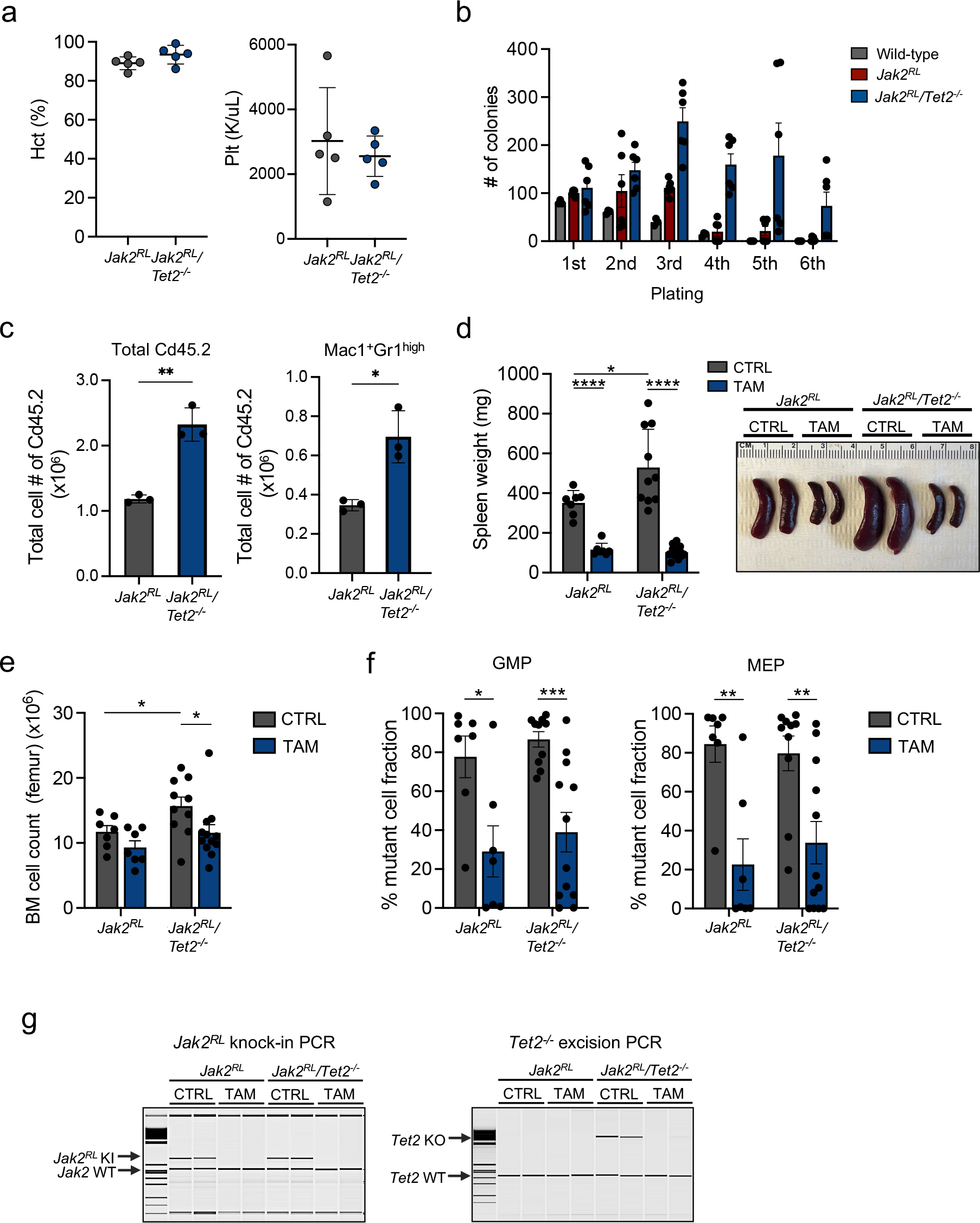
Phenotypic characterization and Jak2^V617F^ oncogenic dependency in the setting of concomitant *Tet2* loss. **a.** Hematocrit (Hct; left panel) and platelet counts (Plt; right panel) of *Jak2^RL^*vs. *Jak2^RL^/Tet2^-/-^* primary transplanted mice at 16 weeks post-transplant (*n* = 5; mean ± s.d.). Representative of an *n* = 2 independent transplants. **b.** Serial replating assay of *Jak2^RL^* vs. *Jak2^RL^/Tet2^-/-^* bone marrow cells in relation to wild-type bone marrow. Colonies were counted at day 8 after each plating (each sample plated in triplicate, *n* = 2 independent experiments, mean ± s.d.). **c.** Total Cd45.2 cell (left panel) and Mac1^+^Gr1^high^ mature neutrophil cell (right panel) output of *Jak2^RL^* vs. *Jak2^RL^/Tet2^-/-^* cells following 7 days of culture over bone marrow endothelial cells (BMECs) (*n* = 3 replicates per arm; mean ± s.d.). Representative of *n* = 2 independent experiments. \**p* ≤ 0.05, \*\**p* ≤ 0.01. **d.** Spleen weights (left panel) and representative isolated spleens (right panel) of MPN (CTRL) vs. tamoxifen (TAM; *Jak2^V617F^*-deleted) treated *Jak2^RL^* vs. *Jak2^RL^/Tet2^-/-^* mice (*n* ≥ 7 biological replicates per arm across 2 independent transplants; mean ± s.d.). \**p* ≤ 0.05, \*\*\*\**p* ≤ 0.0001. **e.** Total bone marrow (BM) cellularity (femur), and **f.** Bone marrow mutant cell fraction within the granulocytic-monocytic progenitor (GMP; Lineage^-^ cKit^+^Sca1^-^Cd34^+^Fcg^+^; left panel) and megakaryocytic-erythroid progenitor (MEP; Lineage^-^ cKit^+^Sca1^-^Cd34^-^Fcg^-^; right panel) compartments of MPN (CTRL) vs. tamoxifen (TAM; *Jak2^V617F^*- deleted) treated *Jak2^RL^* vs. *Jak2^RL^/Tet2^-/-^* mice (*n* ≥ 7 biological replicates per arm across 2 independent transplants; mean ± s.e.m.). \**p* ≤ 0.05, \*\**p* ≤ 0.01, \*\*\**p* ≤ 0.001. **g.** Representative *Jak2^V617F^* knock-in (left panel) and *Tet2^-/-^*excision genotyping (right panel) by polymerase chain reaction (PCR) of genomic DNA isolated from whole bone marrow of MPN (CTRL) vs. tamoxifen (TAM; *Jak2^V617F^*-deleted) treated *Jak2^RL^* vs. *Jak2^RL^/Tet2^-/-^* mice from time of sacrifice demonstrating presence or loss of *Jak2^V617F^*knock-in (KI) and/or *Tet2^-/-^* excised (knock-out; KO) bands in relation to respective wild-type (WT) bands. Representative bands of *n* = 4-7 individual replicates per arm across 2 independent transplants.

